# Crowdsourcing and phylogenetic modelling reveal parrot tool use is not rare

**DOI:** 10.1101/2023.08.14.553302

**Authors:** Amalia P. M. Bastos, Scott Claessens, Ximena J. Nelson, David Welch, Quentin D. Atkinson, Alex H. Taylor

## Abstract

Studying the prevalence of putatively rare behaviours, such as tool use, is challenging because absence of evidence can arise either from a species’ inability to produce the behaviour or from insufficient research effort. Here, we tackle this challenge by combining crowdsourcing and phylogenetic modelling to approximate actual rates of a rarely observed behaviour based on limited data, targeting tool use in parrots. Crowdsourcing on a social media platform revealed novel instances of tool use in 17 parrot species, more than doubling the confirmed number of tool-using parrot species from 11 (3%) to 28 (7%). Phylogenetic modelling ranked additional species that are most likely to be unobserved tool users, suggesting that between 11% and 17% of extant parrot species may be tool users. These discoveries have implications for inferences about the evolutionary drivers and origins of tool use in parrots, revealing associations with relative brain size and feeding generalism and indicating several genera where tool use was likely an ancestral trait. Overall, our findings challenge the assumption that current sampling effort captures the full distribution of putatively rare animal behaviours. Combining our sampling and analysis methods offers a fruitful approach for investigating the distribution, drivers, and origins of other rare behaviours.

This working paper has not yet been peer-reviewed.

---

Crowdsourcing and phylogenetic modelling reveal parrot tool use is not rare Our understanding of the evolution of animal behaviour is built on the assumption that we have access to sufficient data^1–3^. However, this is not always the case. Data on behaviours that are rare, fleeting, or otherwise difficult to observe are likely to be patchy and incomplete^4,5^. Among species for which such behaviours have not been observed, it can be difficult to differentiate between cases in which the species is truly incapable of producing the behaviour and cases in which the species is capable of producing the behaviour but the behaviour has not yet been observed. Such a distinction can be critical for drawing conclusions about the rarity and evolution of the behaviour in question.

Comparative work on the evolution of tool use is a paradigmatic example of this issue. The initial discoveries of tool use in chimpanzees^6^, birds^7^, dolphins^8^, and octopuses^9^ occurred decades after significant advances on other more easily measurable aspects of their biology. Since these initial discoveries, scholars have used the distribution of reported tool use across species to make various claims about the evolutionary drivers of tool use behaviours. For example, based on the observation that bird species with reported tool use tended to have larger brains, researchers have identified higher relative brain size as a likely precondition for tool using capabilities^10–12^. These researchers argue that larger brains are better able to integrate visual and somatosensory information when innovating novel behaviours, such as tool use, in changing environments^13,14^. Similarly, researchers have used existing reports of tool use in birds to debate the roles of generalist versus specialist feeding strategies in driving the evolution of tool use, with some arguing that feeding generalists require technical innovations to expand their dietary niche^13,15,16^ and others arguing that feeding specialists require technical innovations for extractive foraging of specific foods^17,18^.

However, before we can make claims of this kind, we need to know whether current research effort in the literature is sufficient for robust conclusions to be drawn about the evolution of tool use. In fact, evidence suggests that research effort is often systematically biased towards particular taxonomic groups, parts of the world that are easy to access, and species with life history traits that make them easier to study, such as larger distribution ranges and population sizes^19^. This is a crucial limitation because insufficient observation may lead researchers to miss true instances of tool behaviours and thus draw premature conclusions about the evolutionary drivers and origins of tool use. While researchers have attempted to deal with this problem by controlling for the number of scientific papers published on different species, previous work has not yet attempted to quantify and explicitly model the relationship between actual tool-using behaviour and what is reported in the scientific literature. If more tool-using species exist than previously thought, this could have important implications for theories of the evolutionary drivers and origins of tool use and, more generally, for our understanding of how rare this behaviour actually is.

One potentially powerful method for quantifying actual rates of rare animal behaviours is crowdsourcing^20^. In a crowdsourcing study, researchers collect reports from the general public and/or collate and analyse videos posted on social media platforms. This citizen science approach has been widely used in ecology to monitor the distributional patterns of species^21^, but has also recently been used to uncover a variety of rare animal behaviours, including interspecies play in dogs^20^, novel problem-solving behaviours in horses^22^, and socially-learned foraging innovations in cockatoos^23,24^. By casting the net wider than the scientific literature, the crowdsourcing method can provide an indication of the tool-using species that the literature might be missing.

Even after using this crowdsourcing approach, some tool-users could *still* remain unobserved. An additional approach to identify these unobserved species is to specify a causal model of the process that generates the observed data. We propose one such causal model in Figure 1. In this model, we assume that the presence of tool use in the scientific literature (or in crowdsourced reports) is caused by both unobserved tool use capabilities and the number of published studies (or the number of crowdsourced reports) for any given species. Tool users are more likely to be observed if they are well studied, but understudied tool users may go undetected. Furthermore, based on existing theories of the evolution of tool use^10–18^, we propose that the unobserved presence or absence of tool use is additionally caused by relative brain size, feeding strategy, and shared phylogenetic ancestry. Expressing this causal model as a statistical model can suggest further species which are likely to be unobserved tool-users and, simultaneously, test existing theories of the evolutionary drivers of tool use without incorrectly assuming that absence of evidence is evidence of absence.

**Figure 1.**
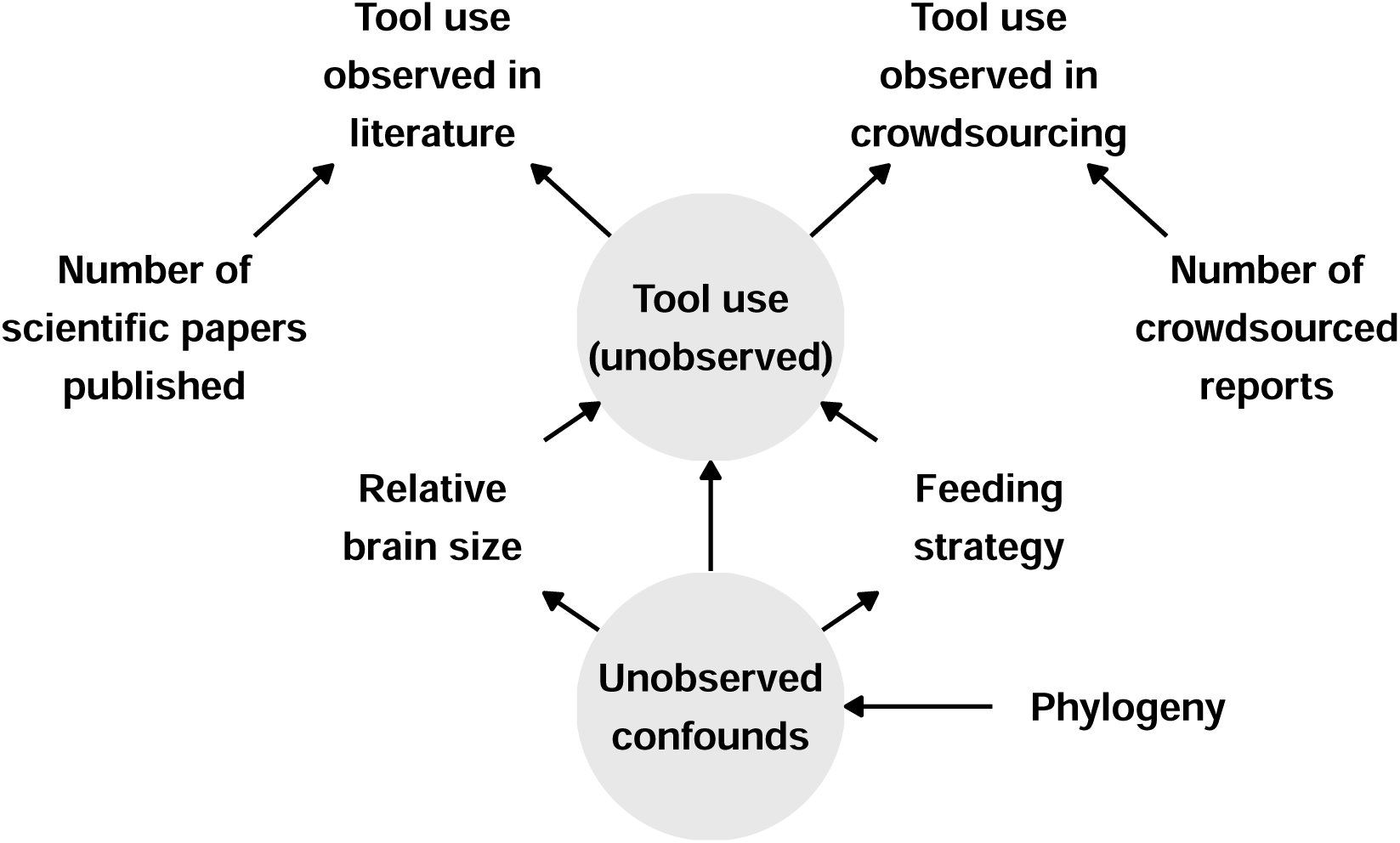
Causal model of observed tool use. Directed acyclic graph of the causal relationships between observed tool use and other variables. Available scientific data on tool use is caused both by unobserved tool use presence and scientific research effort (i.e., number of publications). Available crowdsourced data on tool use is caused both by unobserved tool use presence and crowdsourcing effort (i.e., number of crowdsourced reports). According to theory, unobserved tool use presence should be caused by relative brain size (encephalisation quotient) and feeding strategy (generalist vs. specialist). These variables all share unobserved confounds generated by shared phylogenetic history. Grey circles indicate unobserved variables.

Here, we apply these crowdsourcing and phylogenetic modelling approaches to tool use in the parrot order. We focus on tool use in parrots for a number of reasons. First, the scientific literature suggests that only a small proportion of extant parrot species (11 out of 398^25^; 3%) use tools^10,26–38^ (note that we do not include in this count species that have been previously shown to use touchscreen devices as these behaviours were explicitly trained^39^). Parrot tool use thus provides an ideal test case for examining how robust sampling is in the scientific literature. Second, parrots are highly popular as pets. Over 70% of all extant parrot species are bred in the aviculture industry and kept as pets worldwide^40–46^, enabling us to leverage the power of crowdsourcing on a social media platform to search for evidence of tool use^20^. Third, detailed data on relative brain sizes^47–52^, feeding strategies^53^, and shared ancestry^54^ exist for parrots, allowing us to fit the statistical model implied by Figure 1 to the entire parrot order.

We first present the results from our crowdsourcing survey, in which we collated videos of tool use in parrots from an online video platform. This survey reveals a number of previously unidentified tool-using parrot species, which we map onto the phylogeny of the parrot order. We then describe our statistical model in more detail, and use it to (*i*) rank further parrot species that are likely unobserved tool users and (*ii*) re-examine key hypotheses regarding the evolutionary drivers and origins of tool use in parrots.

## Results

### Crowdsourcing video survey

We surveyed the social media platform YouTube for video evidence of tool use in parrots (see Methods for detailed search criteria). In our search, we defined “true” tool use behaviour as the manipulation of an unattached object to achieve a goal^55^, while “borderline” tool use involved the use of an object that was still attached to a substrate^56^.

In total, we found 116 videos of 104 individuals from 25 parrot species performing either true tool use (100 videos of 89 individuals from 22 species) or borderline tool use (16 videos of 16 individuals from 7 species). All videos featured pet parrots in captive settings. In 68 of these videos, owners did not appear to interfere with the subjects’ actions. In 43 videos, there was potential interference, either from the owners being in close physical contact with the bird (e.g., bird perching on hand), talking to the bird, or handing it the tool (which occurred in only two videos). We could not establish the degree of interference in the remaining 5 videos, as sound had been removed or was substituted by music. None of the videos featured owners directly rewarding tool use behaviours with food. All borderline tool use cases were excluded from further analyses.

Of the 22 parrot species performing true tool use, 13 were represented in our video survey by two or more individuals over multiple independent observations. True tool use always involved the subject using an object for self-scratching (95 videos involved scratching the head and/or neck). The most common tool (53 videos) was a moulted feather. Human-made objects (e.g., pens, spoons, pieces of wood, cardboard) were also common.

According to YouTube video descriptions and owner comments, 45 of the individuals performing true tool use were males and 18 were females. No sex information was provided for the remaining 26 individuals. As owners provided no information on whether sex had been established through genetic testing, and sexual dimorphism in parrots is rare^57,58^, we could not typically ascertain if descriptions were accurate. It is unclear if the disproportionately large number of males in the sample is a consequence of owners more readily assuming their parrots are male when they have not been genetically tested, owners being more likely to own or film male parrots, or male parrots exhibiting more true tool use behaviours than female parrots.

Figure 2 maps the findings from the video survey onto a maximum clade credibility phylogeny for the parrot order, plotted alongside species previously identified in the scientific literature. Before the video survey, 11 parrot species (3%) had been identified as tool users in the scientific literature. Across our video survey, we observed true tool use in 22 species, 5 of which overlapped with the scientific literature and 17 of which were novel species. All of the species identified in the video survey were cockatoos (Cacatuidae), Old World parrots (Psittacinae), or neotropical parrots (Arinae). The most common species in our survey, accounting for 41 videos from 37 individuals, was the green-cheeked conure (*Pyrrhura molinae*). In accordance with the scientific literature, the video survey found no evidence of tool use in any species of Psittaculidae, despite this family containing some of the most commonly kept pet species, including lovebirds, lorikeets, and Asian parakeet species. Combining both the video survey and the scientific literature, we can thus identify 28 tool-using parrot species overall (7%), compared to the 11 previously reported.

**Figure 2.**
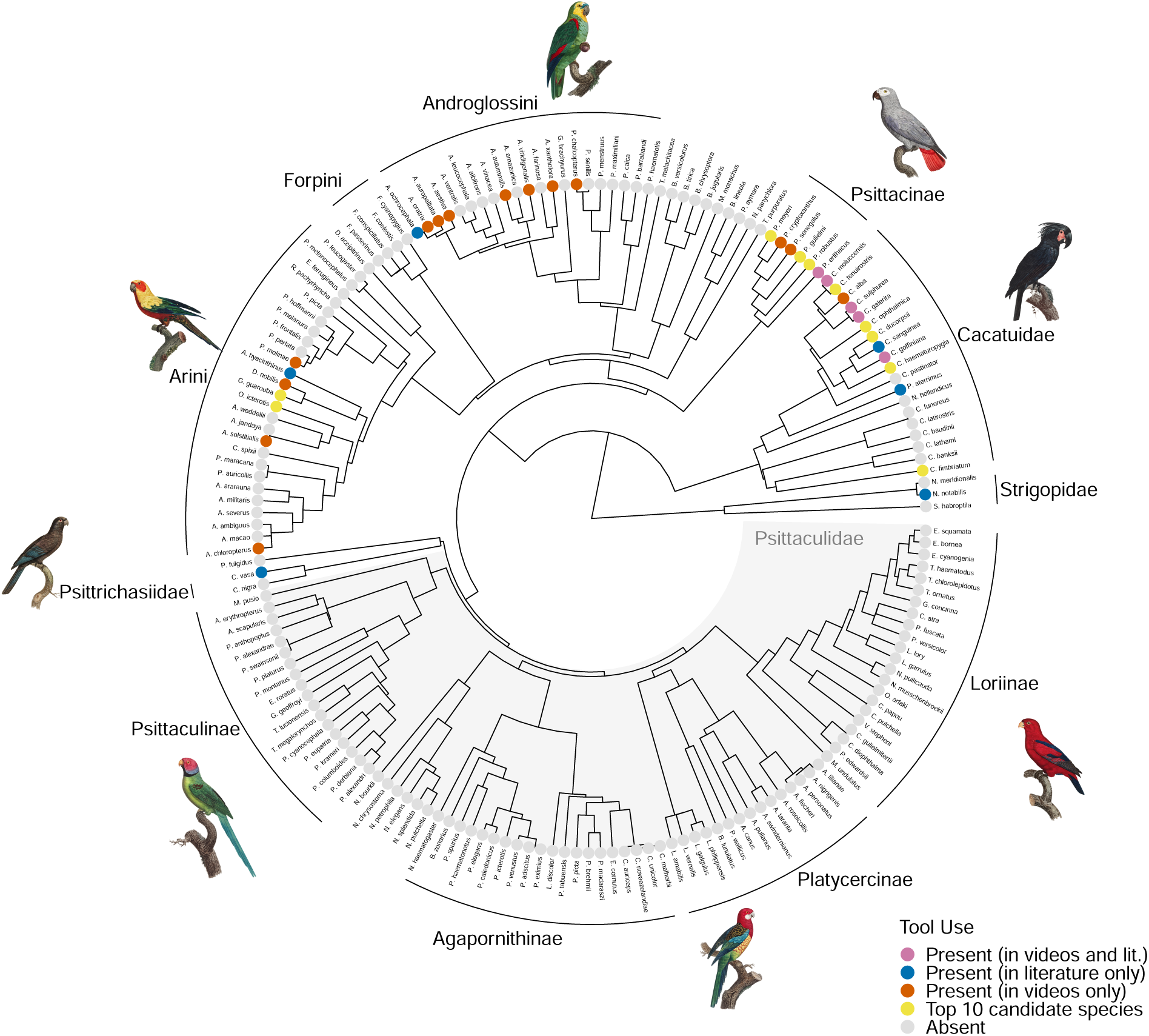
Results of crowdsourcing video survey and phylogenetic survival cure modelling mapped onto a maximum clade credibility phylogeny of the parrot order. Purple tips indicate species observed in the scientific literature and in the video survey. Blue tips indicate species observed only in the scientific literature. Orange tips indicate species observed only in the video survey (note that three species observed only in the video survey are not present in the phylogeny due to a lack of genomic data: *Psittacara erythrogenys*, *Psittacus timneh*, and *Aratinga nenday*). Yellow tips indicate the top ten most likely tool-using species from our phylogenetic survival cure model which were not observed in the scientific literature or the video survey. All other tips are labelled as absent (grey).

The identification of new tool-using species in our video survey increases the extent to which phylogeny can explain the distribution of tool use in the parrot order. We estimated phylogenetic signal (Pagel’s *λ*) of tool use using both the pre-video-survey and post-video-survey data. Pagel’s *λ* varies between 0 and 1, where 0 implies that the distribution of a trait across species is unexplained by phylogenetic relatedness and 1 implies that the distribution of a trait across species is fully explained by phylogeny. Using the evidence of tool use from the scientific literature alone (pre-video-survey data; 11 tool-using species), we estimated an average posterior Pagel’s *λ* of 0.60 (95% credible interval [0.00 0.90]). This estimate was moderate-to-strong, but highly uncertain. In comparison, combining the evidence from both the literature and the video survey (post-video-survey data; 28 tool-using species) resulted in a stronger and more certain estimate of phylogenetic signal. With these data, we estimated Pagel’s *λ* = 0.65 (95% CI [0.50 0.77]). Thus, the results of our video survey increase the extent to which the distribution of tool use across parrot species can be explained by shared phylogenetic ancestry. This suggests that we can potentially use phylogenetic information, along with other variables, to identify further tool-using parrot species that may remain undetected.

### Phylogenetic survival cure model

In addition to the 28 tool-using species identified in the literature and our video survey, we fitted a Bayesian phylogenetic survival cure model to rank further species that are likely to be undetected tool-users (i.e., tool-using species with no tool use reported in the literature or in crowdsourced videos).

Survival cure models are often used in the medical sciences to analyse the time to some event, such as cancer relapse, with censored patient data^59^. The data are right-censored because some patients have relapsed when they are measured, and some have not yet relapsed when they are measured (i.e., they are censored). The cure aspect of these models comes from an additional assumption that a certain proportion of the population is “cured” and can never relapse, no matter how long we measure them for.

Our tool use problem has the same features. We are modelling a time-to-event; specifically, the amount of “time” (i.e., observation opportunities measured as the number of published papers or crowdsourced videos) until tool use is identified. This is right-censored data, because many species will not have had tool use identified when we measure them. Moreover, we can assume that a certain proportion of the population is “cured” – that is, they are not tool users, and so we will never identify tool use for them, no matter how long we measure them for.

In our model, we infer the tool-using status of each species by allowing each species to have their own probability of being “cured” (i.e., not a tool user). Following our causal model (Figure 1), we predict these probabilities based on feeding strategy, encephalisation quotient, and phylogenetic history (see Methods for full model). The model additionally takes research effort into account by allowing that, among species for which tool use is unobserved, all else being equal those with fewer published papers and fewer video search hits have a higher probability of being undetected tool users (Supplementary Figure 1).

We found that this phylogenetic survival cure model was able to adequately distinguish between species with and without evidence for tool use, with an area-under-the-curve classification statistic of 0.95 (Supplementary Figure 2). To further estimate the accuracy of the model’s predictions, we also used a leave-one-species-out approach with known tool users. For each of the 25 tool-using species that were represented on the phylogeny and for which we had brain size and genomic data (we lacked data for three tool-using species), we fitted the model to a modified dataset which set tool use to be absent for the target species in both the scientific literature and the video survey. Across 25 cross-validation models, 18 models (72%) continued to predict the target species as having a median posterior probability of tool use that was within the range of all other tool users. This classification rate was greater than the baseline classification rate of 26% for species without evidence of tool use in the full model (38 of 149 species without evidence of tool use had a median posterior probability of tool use that was within the range of the tool-using species). Together, the area-under-the-curve statistic and the leave-one-species-out approach suggest that the model is able to adequately classify known tool users, with some error.

Figure 3 visualises the ranked posterior probabilities of tool use from the phylogenetic survival cure model for all parrot species. As expected, the known tool users are ranked towards the top of this list. However, several “tool use absent” species also rank highly on the list, despite not being identified as tool users in the scientific literature or in our video survey. In fact, according to the model, the most likely tool user is a species for which tool use is unobserved in our data: the blue-eyed cockatoo (*Cacatua opthalmica*). This species is endemic to Papua New Guinea and is relatively understudied, with only 6 published papers and 596 video search hits, which is fewer than the model expects are necessary to discover tool use when it is present (Figure 4). This species is also found in the *Cacatua* genus, a clade containing several known tool users. This prediction makes sense given the high phylogenetic signal for tool use reported by the model (Supplementary Figures 3 and 4). Beyond the blue-eyed cockatoo, other highly ranked species without observed evidence of tool use are the Meyer’s parrot (*Poicephalus meyeri*), the golden parakeet (*Guaruba guarouba*), the long-billed corella (*Cacatua tenuirostris*), the Solomons cockatoo (*Cacatua ducorpsii*), the red-fronted parrot (*Poicephalus gulielmi*), the Cape parrot (*Poicephalus robustus*), the yellow-eared parrot (*Ognorhynchus icterotis*), the red-vented cockatoo (*Cacatua haematuropygia*), and the gang-gang cockatoo (*Callocephalon fimbriatum*). Figure 2 plots these species on the parrot phylogeny, using the top ten highest ranked species without observed evidence of tool use as an abitrary cutoff for visualisation purposes.

**Figure 3.**
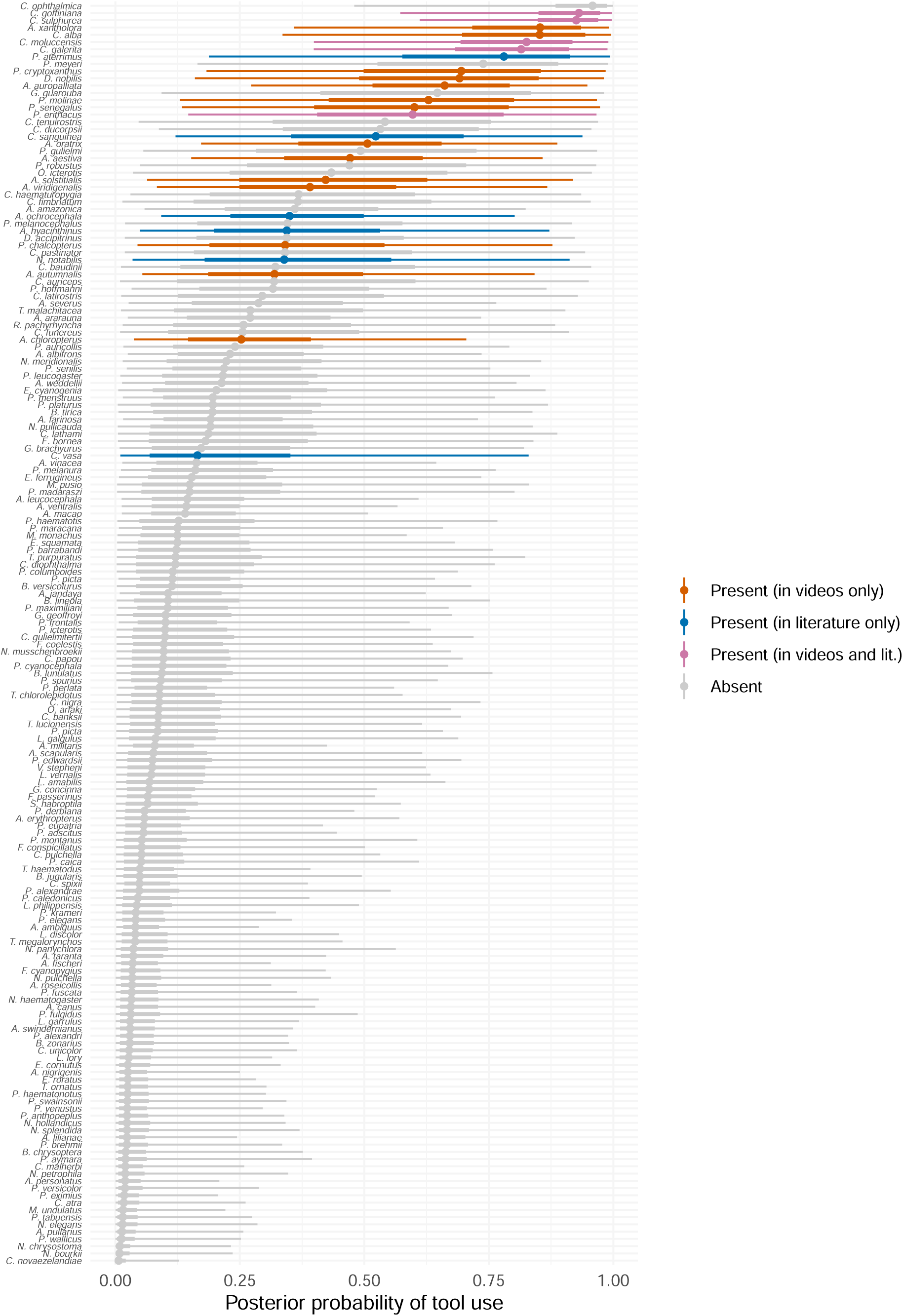
Posterior predicted probabilities of tool use for each species from our phylogenetic survival cure model. Points are posterior medians and lines are 50% and 95% credible intervals.

**Figure 4.**
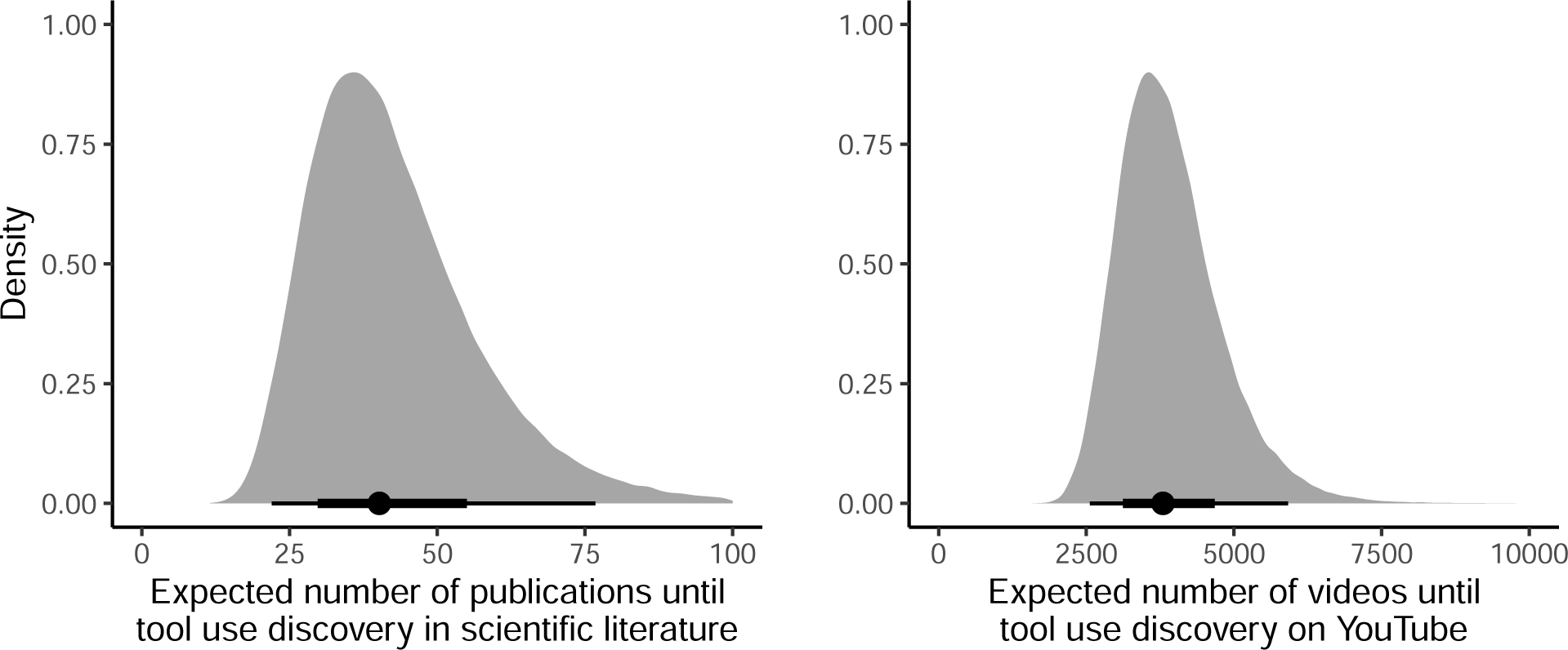
Expected number of published papers and videos until tool use discovery, according to the survival component of the phylogenetic survival cure model. Densities are full posterior distributions, points are posterior medians, and lines are 50% and 95% credible intervals.

The posterior probabilities shown in Figure 3 are estimated with uncertainty, so it is difficult to “identify” any particular species as an undetected tool user. Nevertheless, taking the sum of all the posterior probabilities for the 149 species without observed evidence of tool use, we can estimate that around 26 of those species are likely to be undetected tool users (median sum of probabilities = 25.68, 95% CI [15.15 41.33]). When combined with the species known to use tools, this implies that between 11% and 17% of extant parrot species may be tool users.

### Implications for the evolutionary drivers and origins of tool use

The predicted probabilities from our phylogenetic survival cure model have implications for inferences about the evolutionary drivers and origins of tool use in the parrot order. Regarding the drivers of tool use hypothesised in Figure 1, the phylogenetic survival cure model revealed that encephalisation quotient strongly positively predicted the probability of tool use (median posterior log odds slope = 1.12, 95% CI [0.39 2.00]; Figure 5). This helps explain the ranking in Figure 3: the blue-eyed cockatoo has the largest relative brain size in the dataset. We also found that feeding generalist species were slightly more likely to be tool users, though the posterior difference between generalists and specialists was quite uncertain (median posterior log odds difference = 0.33, 95% CI [-1.13 1.76]). These results from the survival cure model differed from the results of models fitted to pre-video-survey and post-video-survey data without the survival cure component, which found no effect of relative brain size and no difference between feeding strategies, respectively (Supplementary Figure 5).

**Figure 5.**
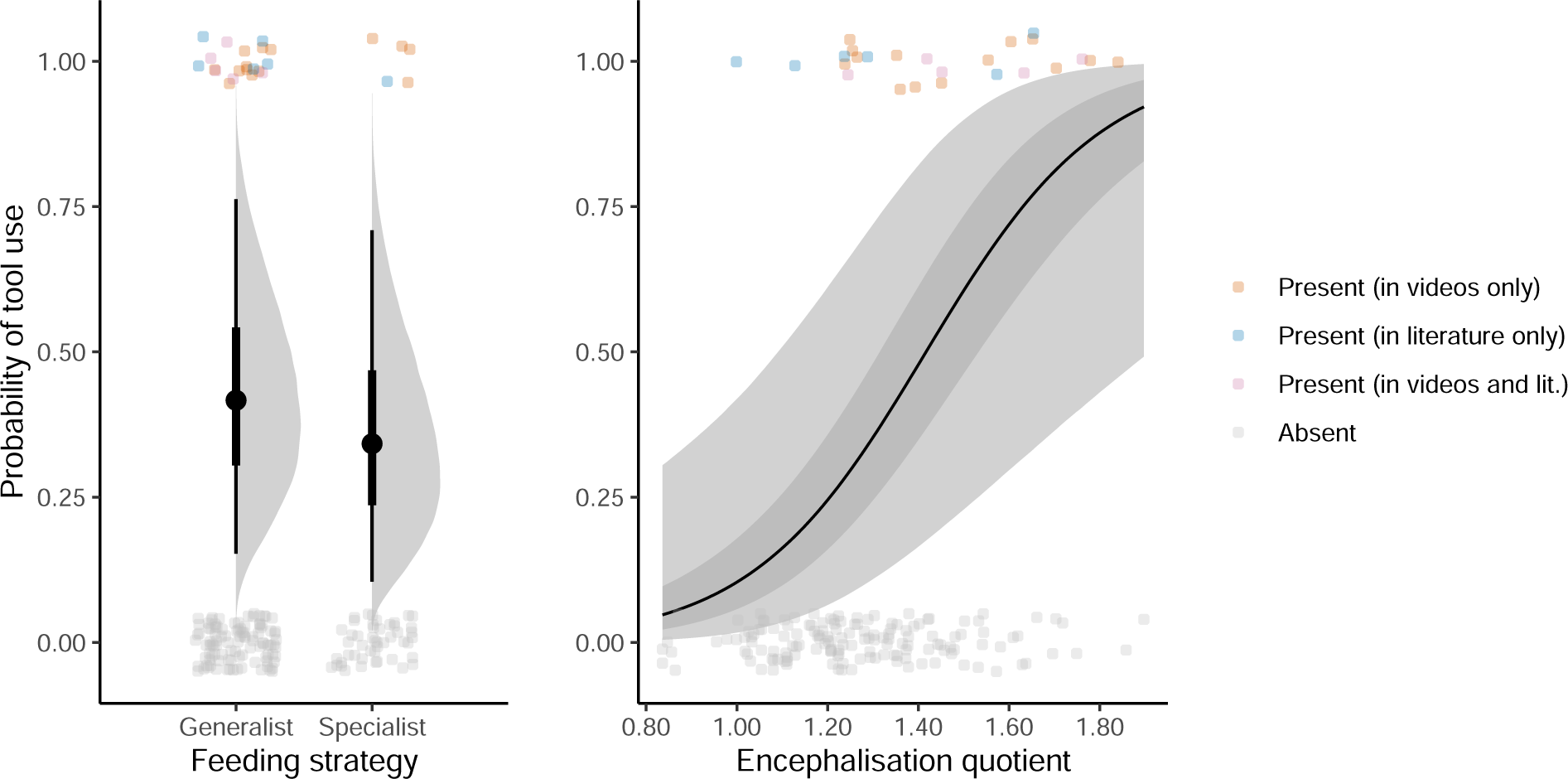
Posterior predictions for the effects of feeding strategy and encephalisation quotient on the probability of tool use from the phylogenetic survival cure model. In the left plot, points and lines represent posterior medians and 50% and 95% credible intervals, with densities representing full posterior distributions. In the right plot, the line and shaded area represents the posterior median regression line with 95% credible intervals. In both plots, individual species are coloured according to the presence / absence of tool use in the video survey and the scientific literature.

Regarding the origins of tool use, we fitted exploratory ancestral state reconstruction models to the pre-video-survey data, the post-video-survey data, and the predicted probabilities from the phylogenetic survival cure model. The discoveries from our video survey and from our phylogenetic modelling increased the likelihood that tool use was present in the most recent common ancestor for several parrot genera. These include the amazon parrots native to the Americas (*Amazona*), the true white cockatoos and corellas found in South East Asia and Australasia (*Cacatua*), the kea and the kākā from New Zealand (*Nestor*), and the *Poicephalus* genus native to Africa (Table 1; Supplementary Figures 6 - 8). These findings suggest that species from each of these genera share their tool use capabilities via common descent from their respective common ancestors, rather than via independent evolution within each genus.

**Table 1.** Estimated probabilities of tool use for most recent common ancestors of several parrot genera. Probabilities estimated using exploratory ancestral state reconstruction models fitted to the pre-video-survey data, post-video-survey data, and predicted probabilities from the phylogenetic survival cure model.

| Genus | Pre-video-survey | Post-video-survey | Survival cure probabilities |
| --- | --- | --- | --- |
| *Amazona* | 0.06, 95% CI [0.01 0.17] | 0.21, 95% CI [0.04 0.66] | 0.48, 95% CI [0.05 0.84] |
| *Cacatua* | 0.08, 95% CI [0.03 0.20] | 0.17, 95% CI [0.03 0.45] | 0.81, 95% CI [0.23 0.99] |
| *Nestor* | 0.11, 95% CI [0.05 0.29] | 0.21, 95% CI [0.12 0.37] | 0.53, 95% CI [0.29 0.66] |
| *Poicephalus* | 0.06, 95% CI [0.02 0.11] | 0.19, 95% CI [0.12 0.41] | 0.72, 95% CI [0.16 0.90] |

## Discussion

Since the earliest anecdotal reports of parrots using tools in the 1970s^60^, only 11 parrot species (3% of all extant parrot species) have been documented as tool users in the scientific literature. Our study used crowdsourcing to identify 17 additional tool-using parrot species that are new to science, more than doubling the overall count to 28 species (7%). These species consisted of cockatoos (Cacatuidae), Old World parrots (Psittacinae), and neotropical parrots (Arinae).

We found strong phylogenetic signal in our extended dataset, demonstrating that the distribution of tool use across parrot species is affected by shared phylogenetic ancestry. This allowed us to use phylogenetic information, along with other variables, to infer the unobserved probabilities of tool use across the parrot order. Our phylogenetic survival cure model incorporated information on phylogenetic history, research effort, relative brain size, and feeding specialisation to rank parrot species that were most likely to be undetected tool users. The sum of probabilities from this model implied that between 15 and 41 of the species without observed evidence of tool use are likely to be undetected tool users, suggesting that the true proportion of tool users may be as high as 17%.

These findings have a number of important implications. First, our findings show that current research effort in the scientific literature is insufficient to capture the real world occurrence of parrot tool use. If the scientific literature had sampled the natural world sufficiently, we would expect to see close correspondence between those species reported as tool users in the literature and those species the public have filmed performing tool use. Instead, we discovered a large discrepancy between these two data sources, both in the prevalence of tool use and the species identified. This raises the possibility that insufficient research effort is a general issue across the scientific literature, both for tool use in other groups and for other rare behaviours.

Second, in terms of the evolution of tool use in parrots, our study challenges a key assumption made in the literature to date: that only a minority of parrots are tool users^30,31,34,37,38^. The paucity of evidence for tool use across parrots in the literature initially implied that tool use may have evolved independently in different parrot species. Our discovery of the widespread distribution of tool use across the parrot phylogeny, along with the strong phylogenetic signal in this expanded dataset, challenges this and suggests that, at least for some parrot clades, the capacity for tool use might be a homologous trait that has been evolutionarily conserved. Our exploratory ancestral state reconstruction analysis provides preliminary support for this hypothesis, revealing an increased probability of tool use among the most recent common ancestors for the *Amazona*, *Cacatua*, *Nestor*, and *Poicephalus* genera. Even at this preliminary stage, our analysis therefore raises an alternative hypothesis for the observed tool use in *Cacatua*^30,38^ and *Nestor* ^26–29,31,34^, namely that tool behaviours have arisen due to the common ancestor being a tool user, rather than from behavioural innovation within a species.

Third, our results support existing theories of the drivers of tool use. We found that encephalisation was strongly positively related to the probability of tool use in our phylogenetic model, supporting previous theories linking relative brain size to increased tool innovation in birds^10–12^ and primates^61^. Nevertheless, we recognise that cross-species correlations between brain size and behaviour are challenging to interpret causally^3,4^, and thus encourage further work on the specific neural correlates of technical intelligence in parrots (see e.g.^62^). In our phylogenetic model, we also found that tool use was somewhat more likely among feeding generalists compared to feeding specialists, although this difference was uncertain. This trend supports previous suggestions that increased cognitive abilities and technical innovation rates are required to expand a generalist species’ dietary niche^15,17,63^. However, the trend contradicts theories linking tool use to dietary specialisation, whereby species eating specific foods that require extractive foraging have higher cognitive ability and are especially prone to using tools (reviewed in ref^17^).

We believe that the observations of tool use in our video survey reflect intentional tool use, rather than accidental object manipulations or trained behaviours. In the crowdsourcing video survey, individual parrots often used tools slowly and repetitively over long periods of time, even across multiple different videos, suggesting that their behaviour was not accidental^64^. Moreover, for 60% of the species we report on, we found two or more videos of repetitive and sustained scratching by different individuals in separate households. Following the assumption that accidental tool use (e.g., briefly holding an object and attempting to scratch oneself simultaneously) is rare^65^, the repeated observation of the same tool use behaviours for scratching by multiple animals of the same species across multiple households suggests that these manipulations are intentional and recurring events rather than rare “accidents” and likely do not represent unusual stereotypies of a single individual. We also found little evidence to suggest that the tool use behaviours were trained or unintentionally cued by the birds’ owners. Over half of the videos coded contained no evidence of human interference with the parrots’ tool use aside from filming the behaviour. Humans only handed parrots their tools in two of the videos, and none of the videos featured owners directly rewarding tool use behaviours with food. Finally, and perhaps most importantly, the high levels of phylogenetic signal in our data provides strong evidence that the observations from our video survey reflect biologically-endowed capacities for tool use rather than accidents or trained behaviours, which would likely appear uniformly across the phylogeny.

While missing data imputation is becoming more common in phylogenetic analyses^66^, the important distinction between absence of evidence and evidence of absence has not been given as much attention. Our phylogenetic analysis provides one approach to this problem by distinguishing between true absences of tool use and absences of tool use due to a lack of research effort in the scientific literature or in crowdsourced videos. To achieve this, we explicitly modelled the measurement of the outcome variable along a research effort time series, such that species with lower research effort in the literature or in videos were likely to be censored. In line with our causal model, we also included relative brain size, feeding strategy, and phylogenetic history as predictors of unobserved tool use. We encourage researchers to test this model by directing future study efforts towards the parrot species with the highest probabilities of being undetected tool-users. Future work should also refine the causal model in Figure 1 to provide more certain estimates of tool use probabilities, either by including additional predictor variables or modelling further causes of measurement error in the taxonomic record, such as species abundance and geographic accessibility^19^.

In conclusion, we have shown that the scientific literature has insufficiently captured the full distribution of tool use in the parrot order. Our crowdsourcing survey has more than doubled the number of known tool using parrot species from 11 to 28, and our phylogenetic inference suggested that the true proportion of parrot tool users could be as high as 17%. These discoveries have implications for theories of the evolutionary drivers and origins of tool use in parrots. Beyond parrot tool use, the crowdsourcing and phylogenetic methods used in this study have the potential to be applied to other rarely observed behaviours, including tool use in other taxa^67^, rhythmic entrainment in birds^68–71^, teasing behaviours in primates^72^, and tactical deception across the animal kingdom^73–75^. We hope that these methods will continue to uncover a diverse array of ephemeral behaviours that have as yet gone undetected in the scientific literature.

## Methods

### Video searches and coding

Our video search was conducted on YouTube in July 2020. Search terms included “parrot using tool” and variants (e.g., “macaw using tool”, “lorikeet using tool”, “parakeet using tool”), “tool use in parrot”, “parrot tool use”, “parrot scratching itself” (included after we found several videos demonstrating self-care tool use in previous searches) and equivalent terms (e.g., “parrot preening itself”, “parrot grooming itself”, “parrot scratching”). For all species that did not display results including object manipulation or scratching behaviours, we also searched the species’ common name(s) + “tool use”, as well as the species’ common name(s) + “scratching”. We also searched for translations of the terms “parrot tool use” and “parrot scratching” in languages for all countries where bird ownership was reported as >5%^76^, namely, Turkish, Czech, Polish, French, Italian, Dutch, German, Russian, Spanish, Portuguese, and Mandarin.

When we found a relevant video, we also searched for similar content uploaded by the same person/channel. For each YouTube search conducted, we watched all relevant videos until we reached five consecutive videos that did not feature any parrots. At this point, we ended that search and initiated the next search. In line with previous recommendations^20^, we planned to exclude any videos that consisted of four or more shots edited together so as to ensure the behaviours being observed were not edited or manipulated, but none of the videos obtained qualified for exclusion.

All videos featuring parrots manipulating objects were investigated for potential tool use or borderline tool use. We defined tool use as the manipulation of an unattached object as an extension of the beak or foot to achieve a goal towards another object, individual, or oneself^55^. Borderline tool use was similarly defined, except that it involved the use of an object that was still attached to a substrate^56^. For example, if individuals used a fallen feather or stick for self-scratching this was defined as tool use, but using one’s currently attached tail feathers or cage furnishings for the same purpose was defined as borderline tool use. Self-scratching had to involve slow and repeated movements of touching an object to one’s body (or, in the case of borderline tool use, rubbing repetitively against an attached object^64^).

All relevant videos were coded for video length, species, tool use presence (yes/borderline), tool use type (e.g., scratching, feeding), the object being used (e.g., feather, stick), tool use target, human interference with the action (talking or handing object to parrot, holding parrot), and the number of shots within each video. Our complete dataset also includes the name for each video, link, subject name, sex (as declared by owner, as most parrot species are not sexually dimorphic), publishing date, and dates found and coded.

### Data for parrot species

We collected data for 194 parrot species (Supplementary Figures 9 - 11). We gathered feeding strategy data as a dichotomous variable (“generalist” or “specialist”) from the EltonTraits ecological database^53^. As per the database, specialists were defined as species whose diet comprised at least 70% of a single food source. To calculate relative brain size, we collated data from the literature for all known body mass (g) and brain mass (g) values across parrots^47–52^. For all species for which we obtained body and brain mass data, we calculated the encephalisation quotient (EQ) using the following formula^77^: *BrainW eight/*(0.12 *∗ BodyW eight* (^2^)). We found body mass and brain mass data for a total of 194 parrot species. This included all tool-using species in our video dataset, with the exception of three species: *Diopsittaca nobilis*, *Psittacara erythrogenys*, and *Coracopsis vasa*. For the latter, we used values for the closely related *Coracopsis nigra*. The other two species were excluded from the final dataset.

For modelling purposes, we coded research effort in both the scientific literature and the crowdsourced videos. For the scientific literature, we operationalised research effort as the number of papers published for each species’ Latin name up to and including the first paper containing tool use for that species. If no tool use had been identified in the scientific literature for a species, then we coded the total number of papers published to date. We used the scientific database Scopus for coding the number of published papers. For the crowdsourced videos, we coded research effort as the number of search hits for each species on YouTube. If tool use had been identified on YouTube, we estimated the number of search hits when the first video of tool use was published on YouTube, assuming linear growth of search hits since the inception of YouTube. If tool use had not been identified, we used the current number of search hits.

For phylogenetic data, we used the phylogenetic tool at www.birdtree.org^54^ to compile 1000 posterior draws of phylogenetic trees for 174 of the 194 parrot species for which both EQ and genomic data exist. A single maximum clade credibility tree was generated from these posterior draws for visualisation purposes. In our analyses, we iterated over posterior draws of the phylogeny to account for phylogenetic uncertainty.

### Phylogenetic signal

We used the *fitDiscrete* function in the *ape* R package^78^ to calculate phylogenetic signal, for both the pre-survey and post-survey tool use data. We iterated the model over 100 posterior parrot phylogenies to incorporate phylogenetic uncertainty.

### Causal model of tool use

To infer unobserved probabilities of tool use across parrots, we proposed a causal model of observed tool use (Figure 1). We assumed that observed tool use in the scientific literature and in the crowdsourced videos is caused by both the unobserved presence or absence of tool use and research effort, proxied by the number of papers published on a species and the number of videos published on a species. Tool users are more likely to be observed if they are well studied, but understudied tool users may go undetected. In addition, based on theory, we also assumed that unobserved tool use is caused by feeding strategy and relative brain size^10–13,15–17^. Finally, we assumed that shared phylogenetic history causes unobserved confounding and non-independence in unobserved tool use, feeding strategy, and relative brain size across the parrot phylogeny.

### Bayesian phylogenetic survival cure model

Given our proposed causal model, we constructed a statistical model to impute unobserved probabilities of tool use and test existing theories of the evolution of tool use in parrots. To understand the model, suppose that we have the following observed variables for parrot species *i*. For the scientific literature, we declare N_Lit_*_,i_* as the number of papers published before and up to tool use identification for species *i* (or, if tool use has not been identified, the total number of papers published for species *i*) and T_Lit_*_,i_* as a binary variable stating whether (1) or not (0) tool use has yet been observed in the scientific literature for species *i*. For the crowdsourced videos, we declare N_Vid_*_,i_* as the number of videos published before and up to tool use identification for species *i* (or, if tool use has not been identified, the total number of videos published for species *i*) and T_Vid_*_,i_* as a binary variable stating whether (1) or not (0) tool use has yet been observed in the crowdsourced videos for species *i*. Additionally, F*_i_* and EQ*_i_* are feeding strategy and encephalisation quotient values for species *i* and we have a phylogenetic distance matrix *D* that describes the patristic distances between all parrot species.

We assume that species *i* is a non-tool-user (i.e. “cured”) with some probability *p_i_*. We also assume that tool use is identified in the scientific literature and the crowdsourced videos at constant rates *λ*_Lit_ and *λ*_Vid_ following exponential survival functions. Given these assumptions, we can then describe the different ways in which variables N_Lit_ and N_Vid_ can be distributed. Focusing on the scientific literature, if tool use has been observed (T_Lit_*_,i_* = 1), then the likelihood for N_Lit_*_,i_* is:

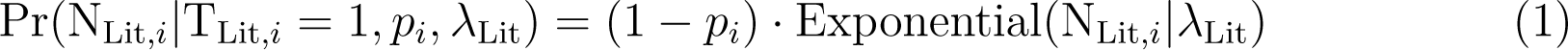

On the other hand, if tool use has not yet been observed (T_Lit_*_,i_* = 0), there are two ways that the outcome variable could have been realised. First, the species could be a non-tool-user with probability *p_i_*. Second, the species could be a tool-user with probability (1 *− p_i_*) that has been censored and has not had its tool use measured yet. Together, then, the likelihood for N_Lit_*_,i_* is:

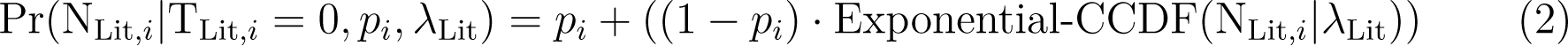

The Exponential-CCDF function allows for the censored nature of the data. The same data generating process is assumed to underlie the crowdsourced videos.

We define the mixture likelihood SurvivalCure as the distribution above, with parameters *p* (the probability of being a “cured” non-tool-user) and *λ* (the rate of the exponential distribution). We use an Ornstein-Uhlenbeck Gaussian process^79^ to model phylogenetic covariance. Below, we specify the full model with priors:

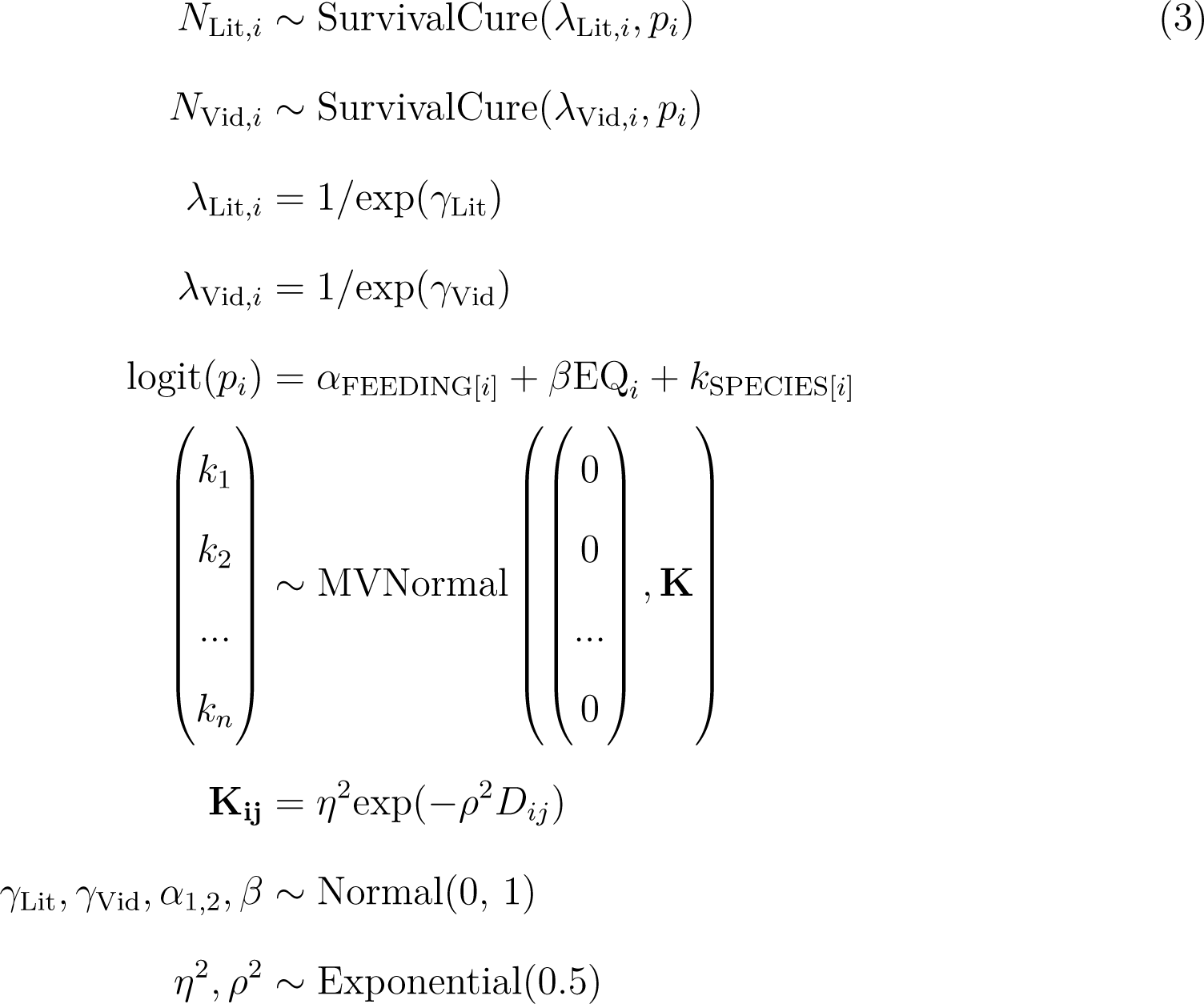

The priors for this model produce reasonable prior predictions of the probabilities of tool use for each parrot species (Supplementary Figure 12), but a sensitivity analysis revealed that the ranking and posterior probabilities reported in the main text were robust to modifying these priors (Supplementary Figure 13). We estimated the posterior distribution of this model using Hamiltonian Monte Carlo as implemented in Stan version 2.26.1^80^. We iterated the model over 100 posterior parrot phylogenies to incorporate phylogenetic uncertainty. R-hat values and effective sample sizes suggested that the model converged normally. Trace plots are reported in Supplementary Figure 14.

To validate our method, we fitted the model to 100 simulated datasets with known parameters. The model was able to successfully recover those parameters (Supplementary Figure 15). We also ran a leave-one-species-out exercise to ensure that we could accurately predict known tool users. We repeated this approach for each known tool user by setting observed tool use to zero. Cross-validation results are reported in the main text.

### Ancestral state reconstruction

To determine whether the identification of novel tool-using species has implications for our understanding of the evolutionary origins of tool use in parrots, we fitted three exploratory ancestral state reconstruction models. We used the *ancThresh* function from the *phytools* R package^81^, iterating the function over 100 posterior parrot phylogenies. This function estimates discrete ancestral states by assuming the evolution of a latent continuous variable following an Ornstein-Uhlenbeck process. We fitted this model to three different outcome variables: (*i*) presence vs. absence of tool use in scientific literature only, (*ii*) presence vs. absence of tool use in literature and/or videos, and (*iii*) the median predicted probabilities of tool use from the phylogenetic survival cure model.

### Reproducibility

All analyses were conducted in R v4.2.1.^82^. Visualisations were produced using the *ggplot2* ^83^ and *cowplot*^84^ packages. The manuscript was reproducibly generated using the *targets*^85^ and *papaja*^86^ packages. Code to reproduce all analyses and figures can be found here: https://github.com/ScottClaessens/phyloParrot

## Acknowledgements

This project was made possible through the support of a grant from the Templeton World Charity Foundation (A.H.T., X.J.N.) and a Prime Minister’s McDiarmid Emerging Scientist Prize (A.H.T.). The authors would like to thank Daniel Sol for providing feedback on a previous version of the manuscript.

## Author Contributions

All authors contributed to the conceptualisation of the paper. A.P.M.B., X.J.N., and A.H.T. developed the video search methodology. S.C., D.W., and Q.A.D. developed the statistical models and analysed the data. All the authors wrote the manuscript and approved the final version for submission.

## Data Availability

All data used in this study are publicly available on GitHub: https://github.com/ScottClaessens/phyloParrot

## Code Availability

All code to reproduce the analyses in this study are publicly available on GitHub: https://github.com/ScottClaessens/phyloParrot

