## Supplementary Information for "Crowdsourcing and phylogenetic modelling reveal parrot tool use is not rare"

### Supplementary Figures

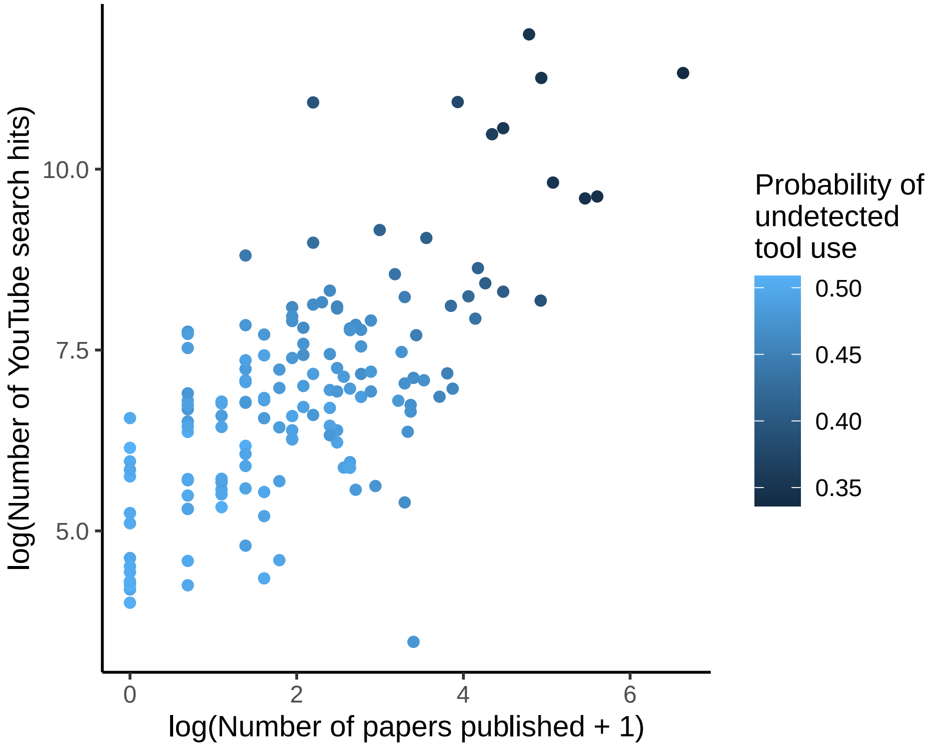


*Supplementary Figure 1. Median posterior probabilities of undetected tool use for each parrot species without observed evidence of tool use from reduced model.* This reduced version of the phylogenetic survival cure model does not contain relative brain size or feeding strategy as predictors, nor does it contain any phylogenetic covariance. The only information included in the model is the number of papers published and the number of YouTube search hits for each species. Each point is a parrot species without observed evidence of tool use, and the colour of the points scales with the probability of undetected tool use. All else being equal, those species with fewer published papers and fewer YouTube search hits have a higher probability of being undetected tool users.

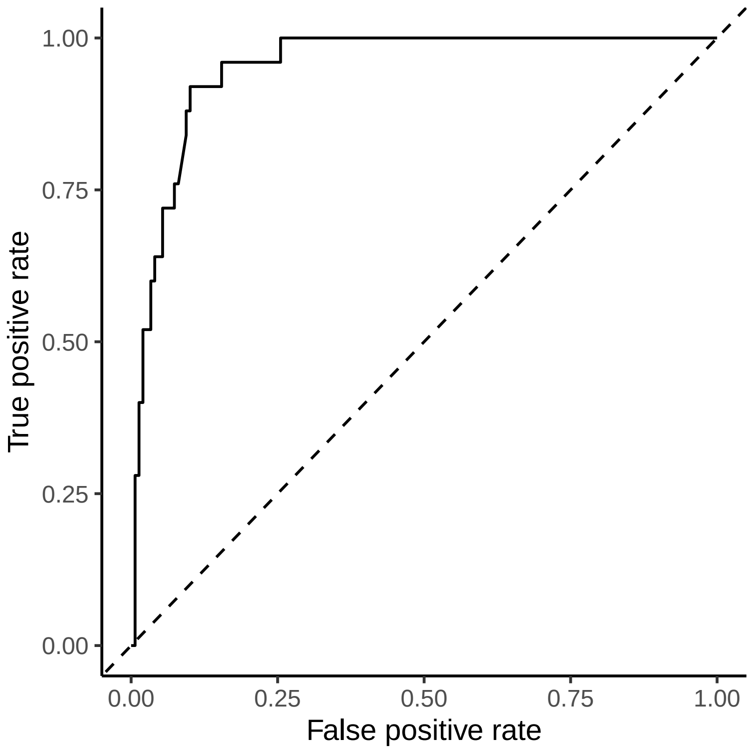


*Supplementary Figure 2. Receiver operating characteristic (ROC) curve for the phylogenetic survival cure model.* The area-under-the-curve in this plot is 0.95, suggesting that the model is able to adequately classify observed tool users and non-tool users.

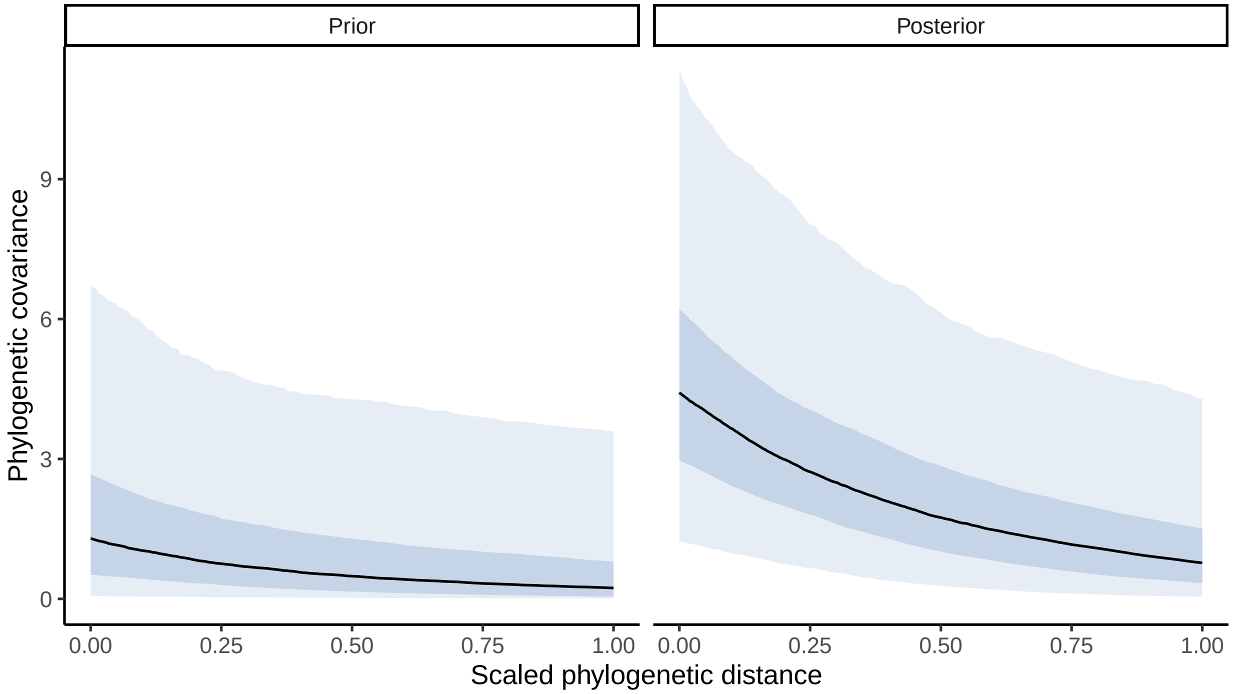


*Supplementary Figure 3*. *Prior and posterior phylogenetic covariance functions from the Bayesian survival cure model fitted to the full dataset.* Lines are median posterior functions and shaded areas are 50% and 95% credible intervals.

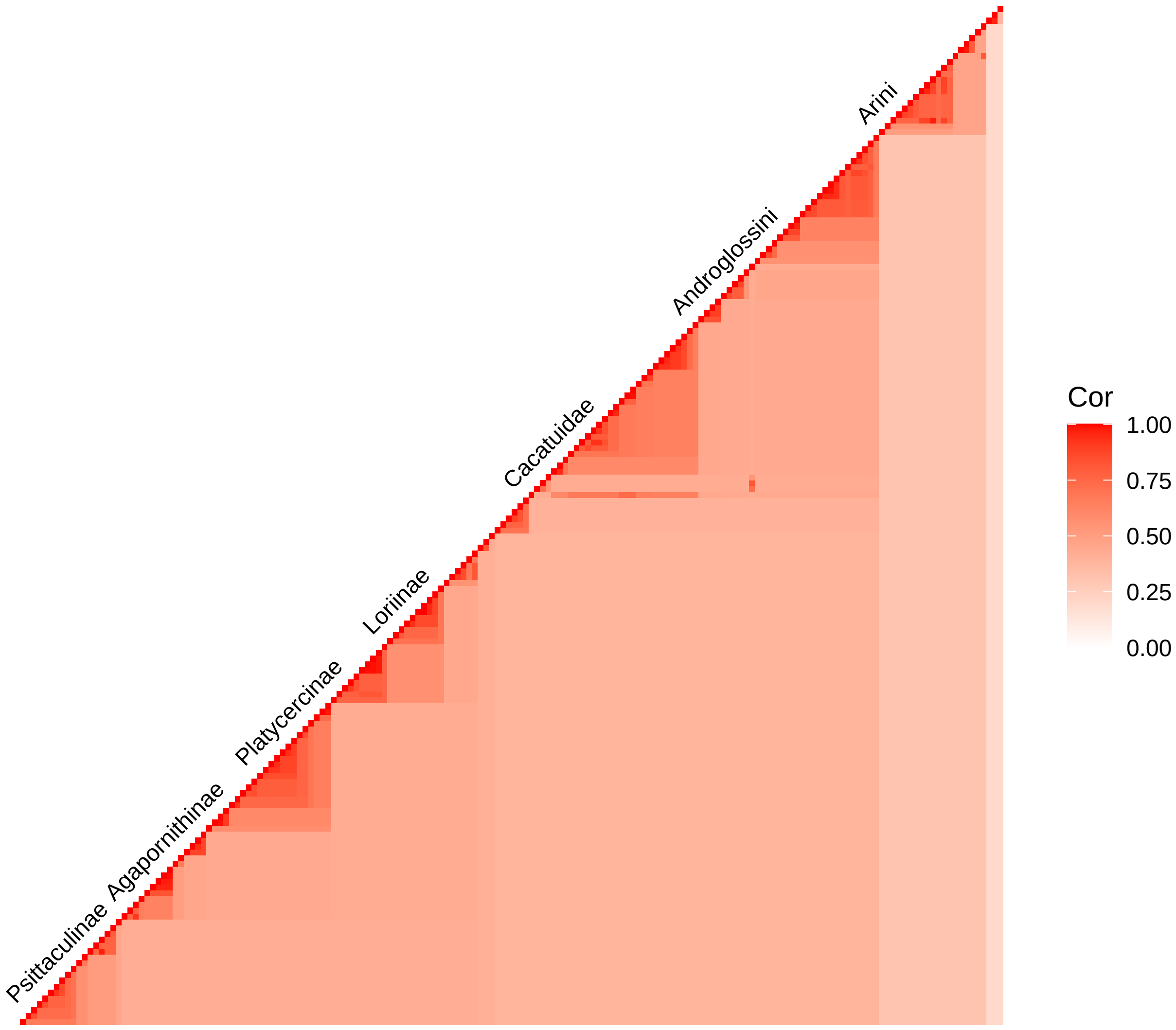


*Supplementary Figure 4. Between-species correlation matrix implied by the posterior phylogenetic covariance function from the Bayesian survival cure model.* Correlations are median posterior estimates. Individual species names omitted for space reasons.

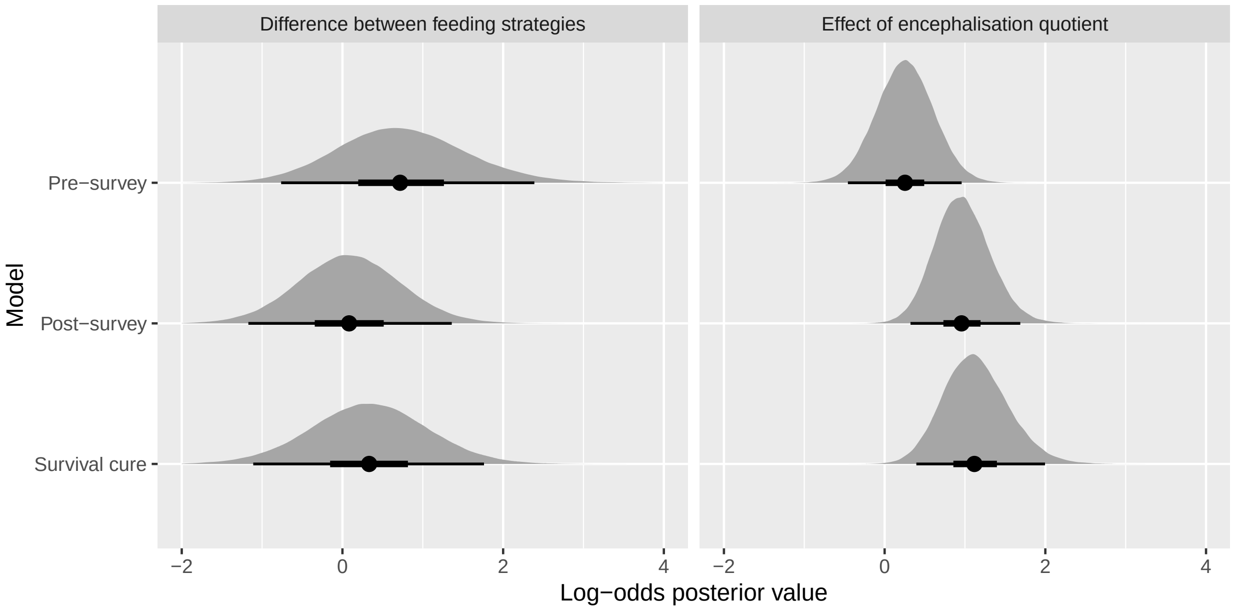


*Supplementary Figure 5. Comparing results between the survival cure model and models fitted to the pre-survey and post-survey data without any survival cure component.* Densities are full posterior distributions from three separate models iterated over 100 posterior parrot phylogenies. Points represent posterior medians, and lines represent 50% and 95% credible intervals.

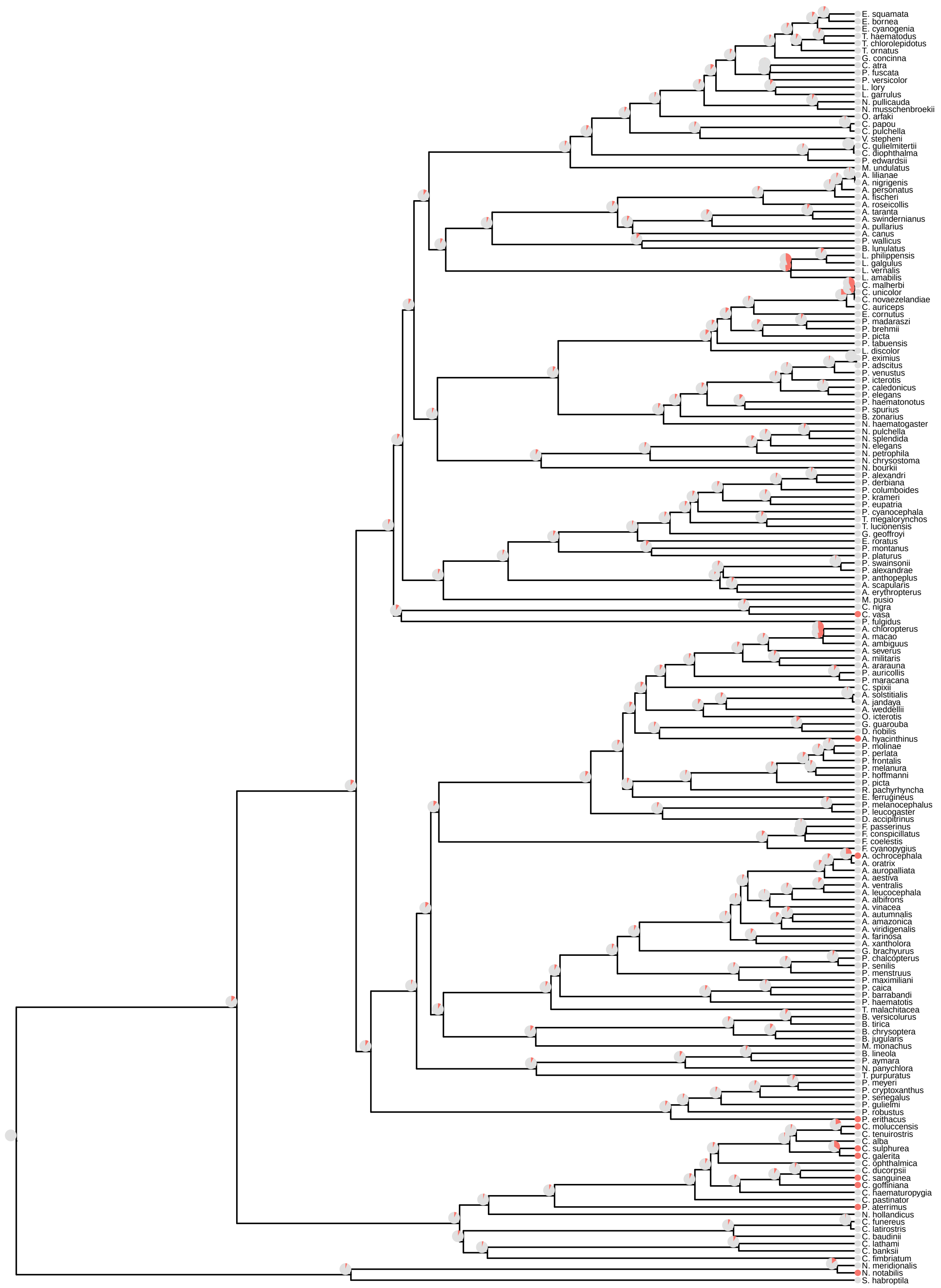


*Supplementary Figure 6. Results of exploratory ancestral state reconstruction analysis fitted to pre-video-survey data, represented on a maximum clade credibility tree.* Tip nodes represent the presence (red) or absence (grey) of observed tool use in the scientific literature. Pie charts represent the posterior probability of tool use presence at each ancestral node.

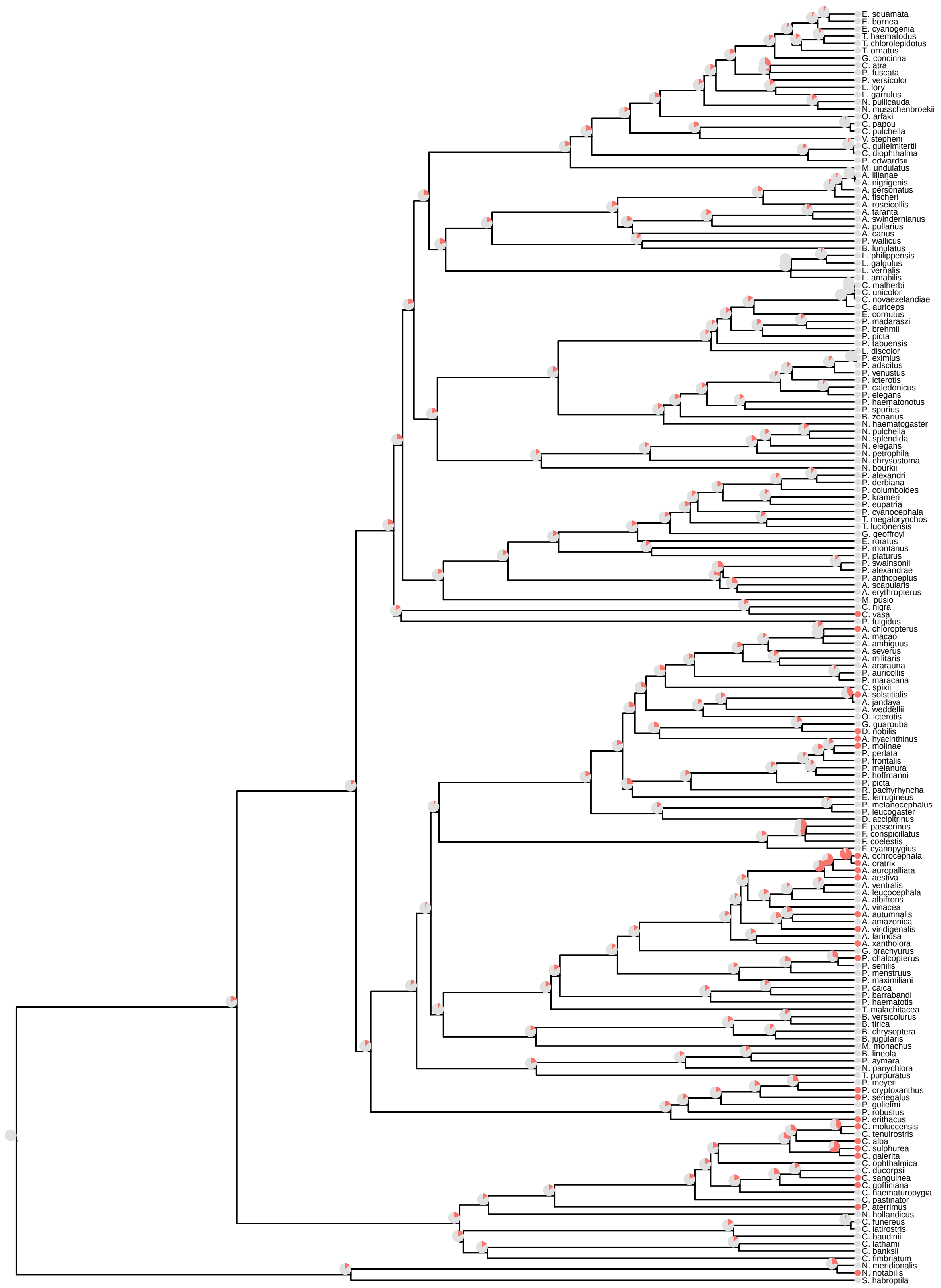


*Supplementary Figure 7. Results of exploratory ancestral state reconstruction analysis fitted to post-video-survey data, represented on a maximum clade credibility tree.* Tip nodes represent the presence (red) or absence (grey) of observed tool use in the scientific literature and the video survey. Pie charts represent the posterior probability of tool use presence at each ancestral node.

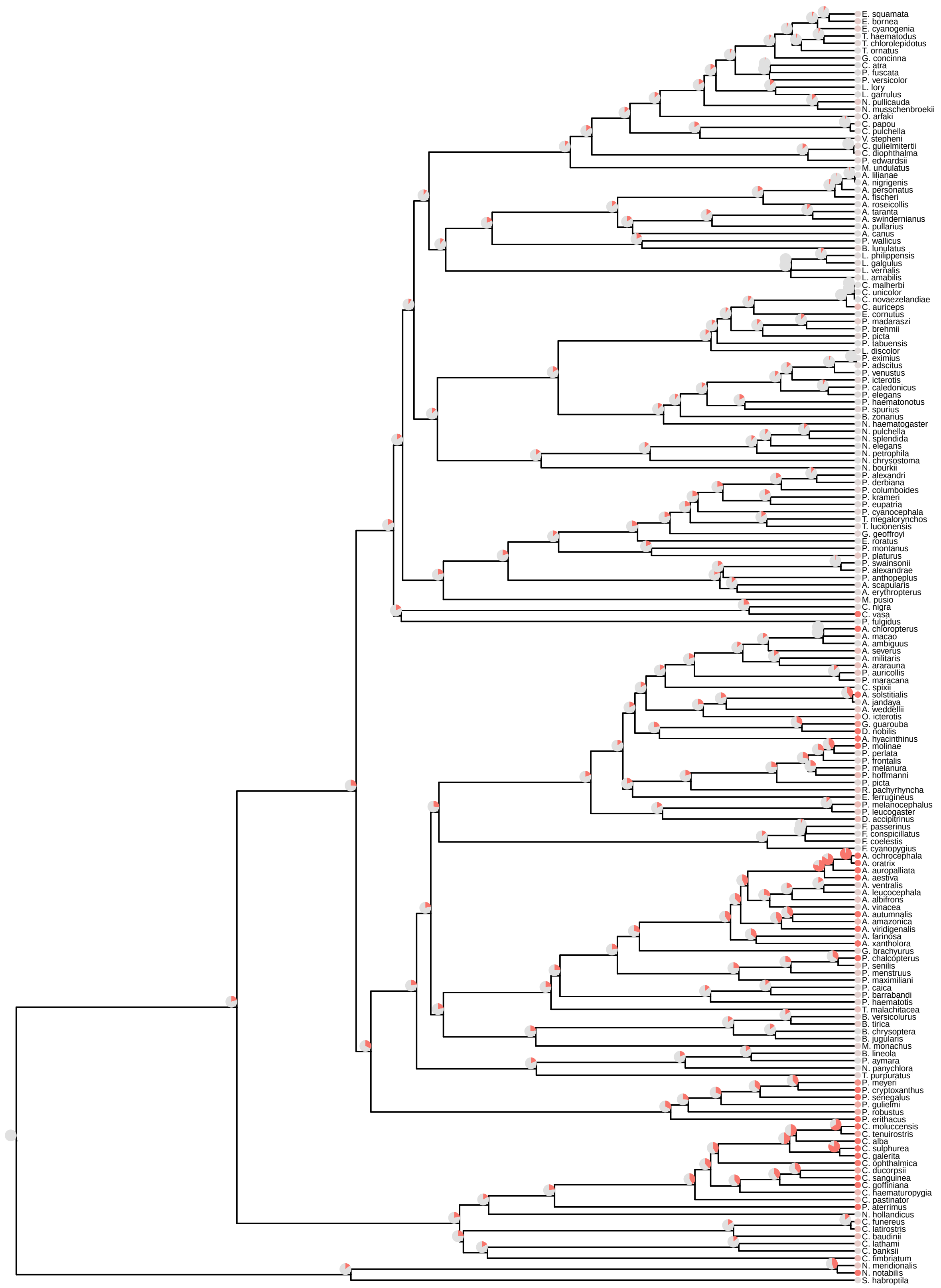


*Supplementary Figure 8. Results of exploratory ancestral state reconstruction analysis fitted to predicted probabilities from the phylogenetic survival cure model, represented on a maximum clade credibility tree.* Tip nodes represent the median posterior predicted probabilities of tool use from the phylogenetic survival cure model, with more red indicating an increasing probability of tool use presence and more grey indicating a decreasing probability of tool use presence. Pie charts represent the posterior probability of tool use presence at each ancestral node.

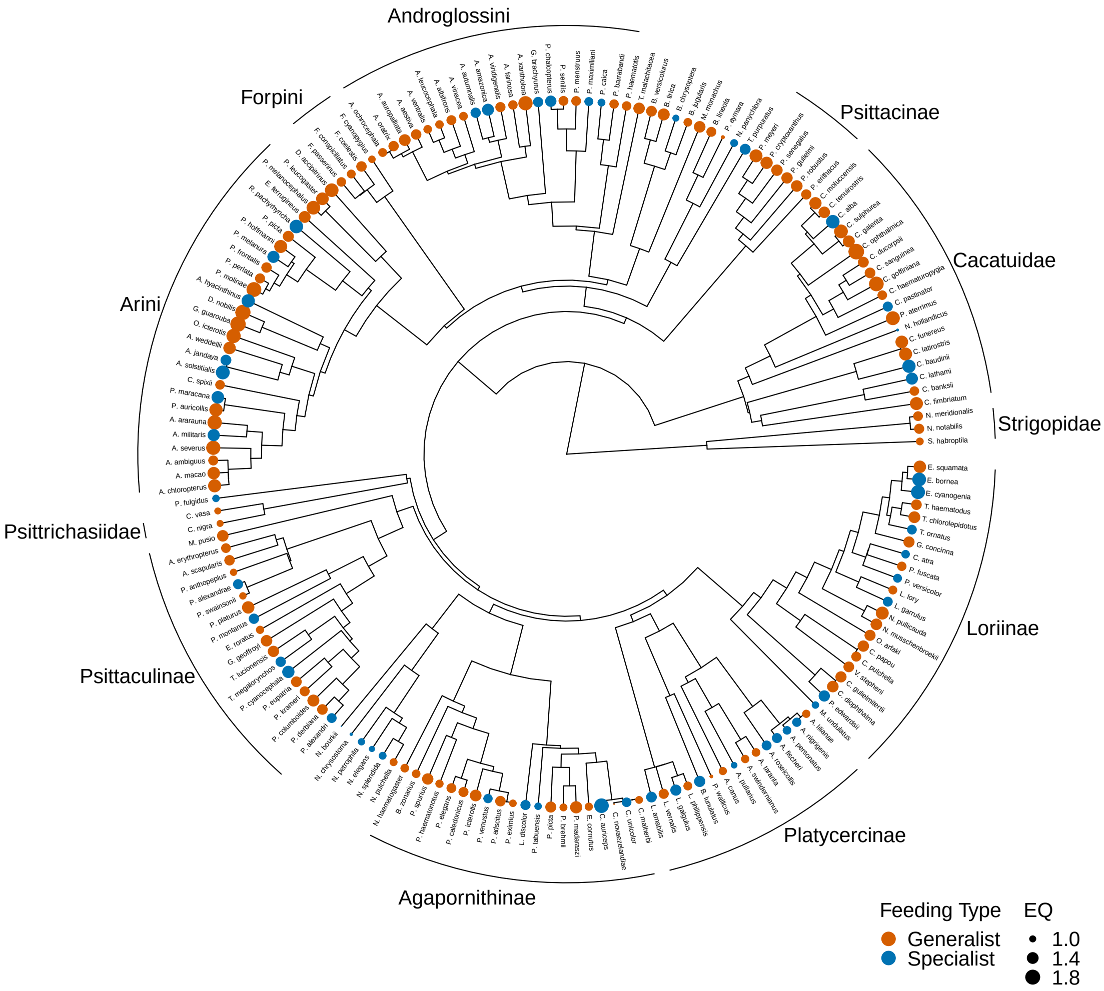


*Supplementary Figure 9. Data on encephalisation quotient and feeding strategy for all parrots, presented on a maximum clade credibility tree.* Tip points are coloured according to feeding generalism (orange) and specialism (blue), and scaled according to encephalisation quotient (EQ).

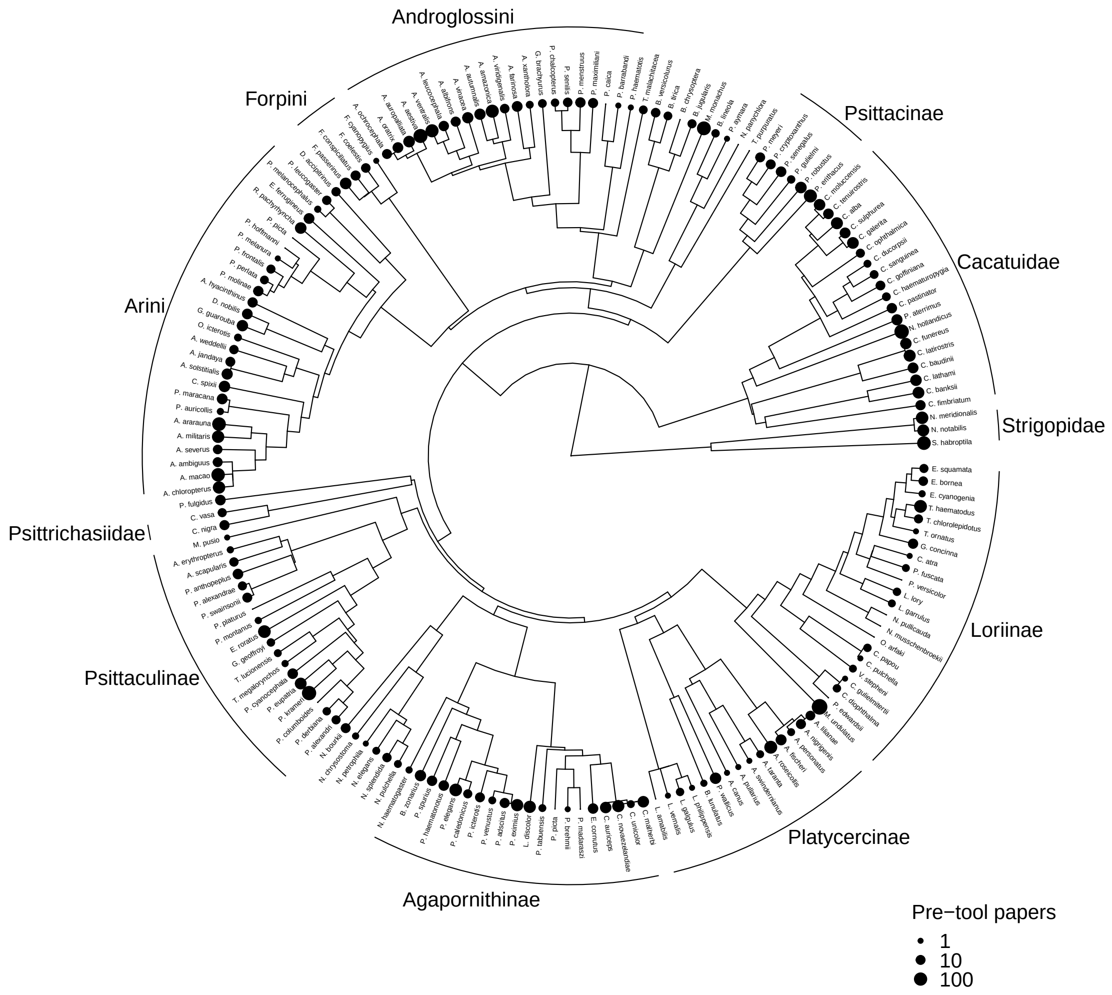


*Supplementary Figure 10. Data on number of scientific publications until tool use discovery for all parrots, presented on a maximum clade credibility tree.* Tip points are scaled according to the number of published papers up until tool use discovery (or, if tool use has not been observed, the current number of published papers).

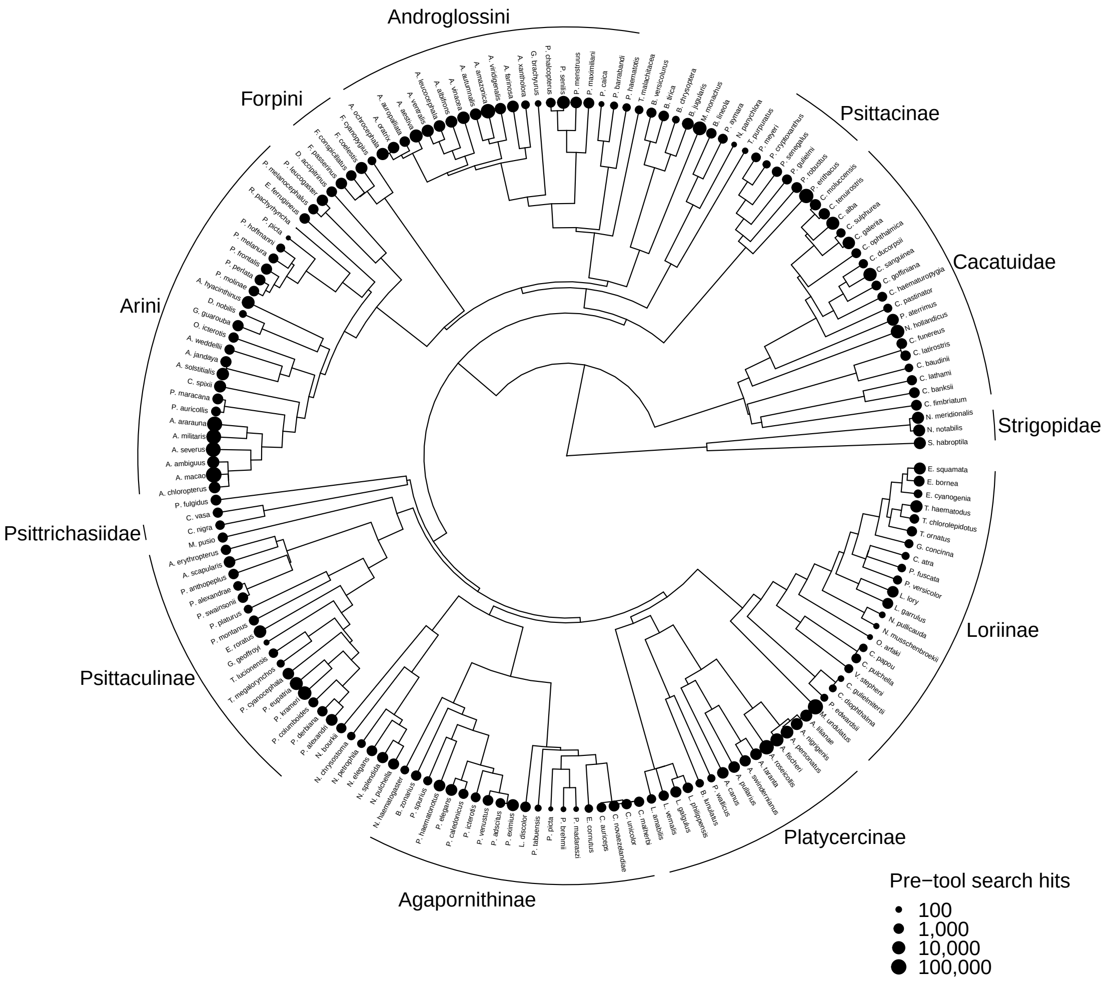


*Supplementary Figure 11. Data on number of video search hits until tool use discovery for all parrots, presented on a maximum clade credibility tree.* Tip points are scaled according to the estimated number of video search hits up until tool use discovery (or, if tool use has not been observed, the current number of video search hits).

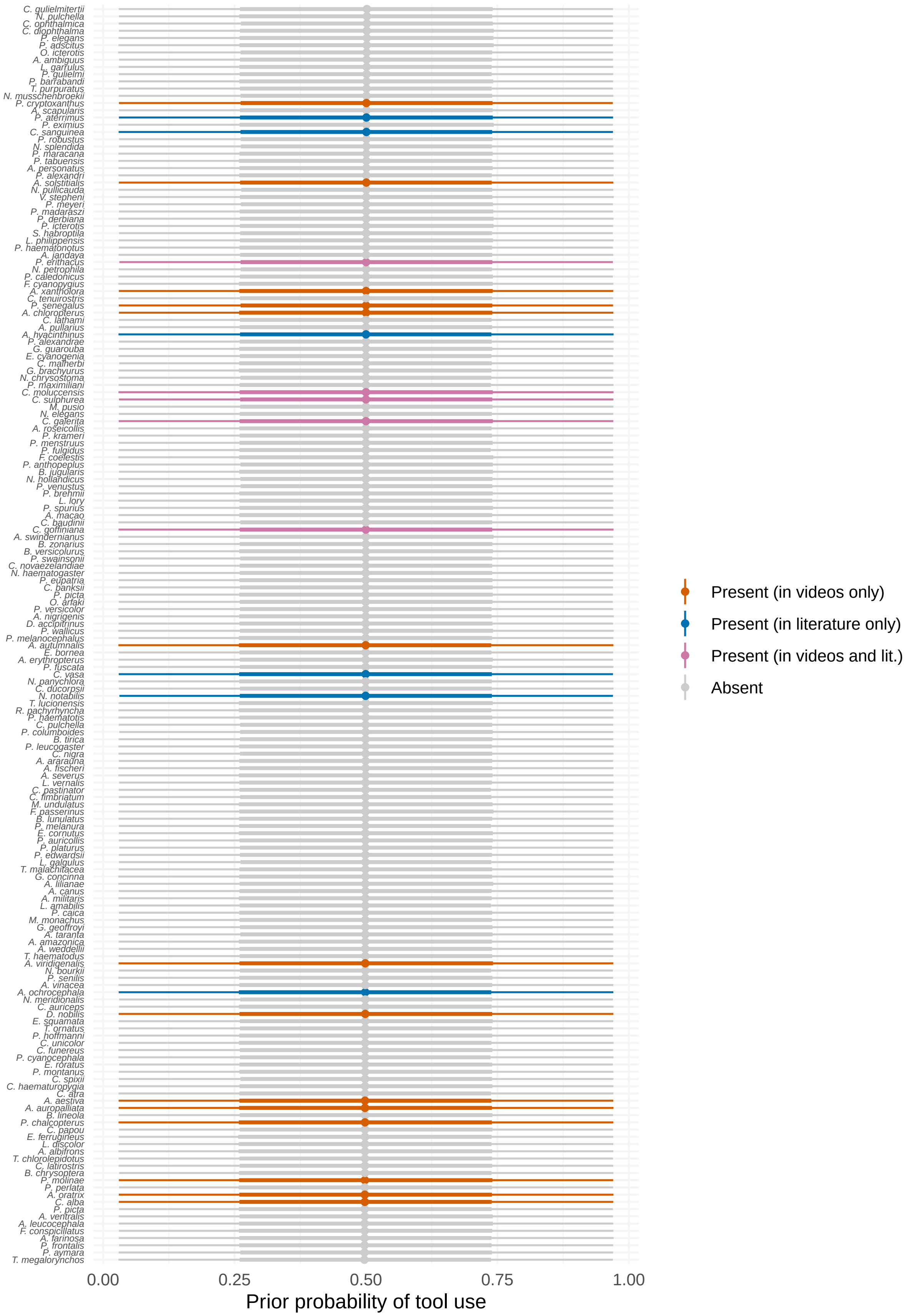


*Supplementary Figure 12. Prior predicted probabilities of tool use for each species from our phylogenetic survival cure model.* Points are prior medians and lines are 50% and 95% credible intervals.

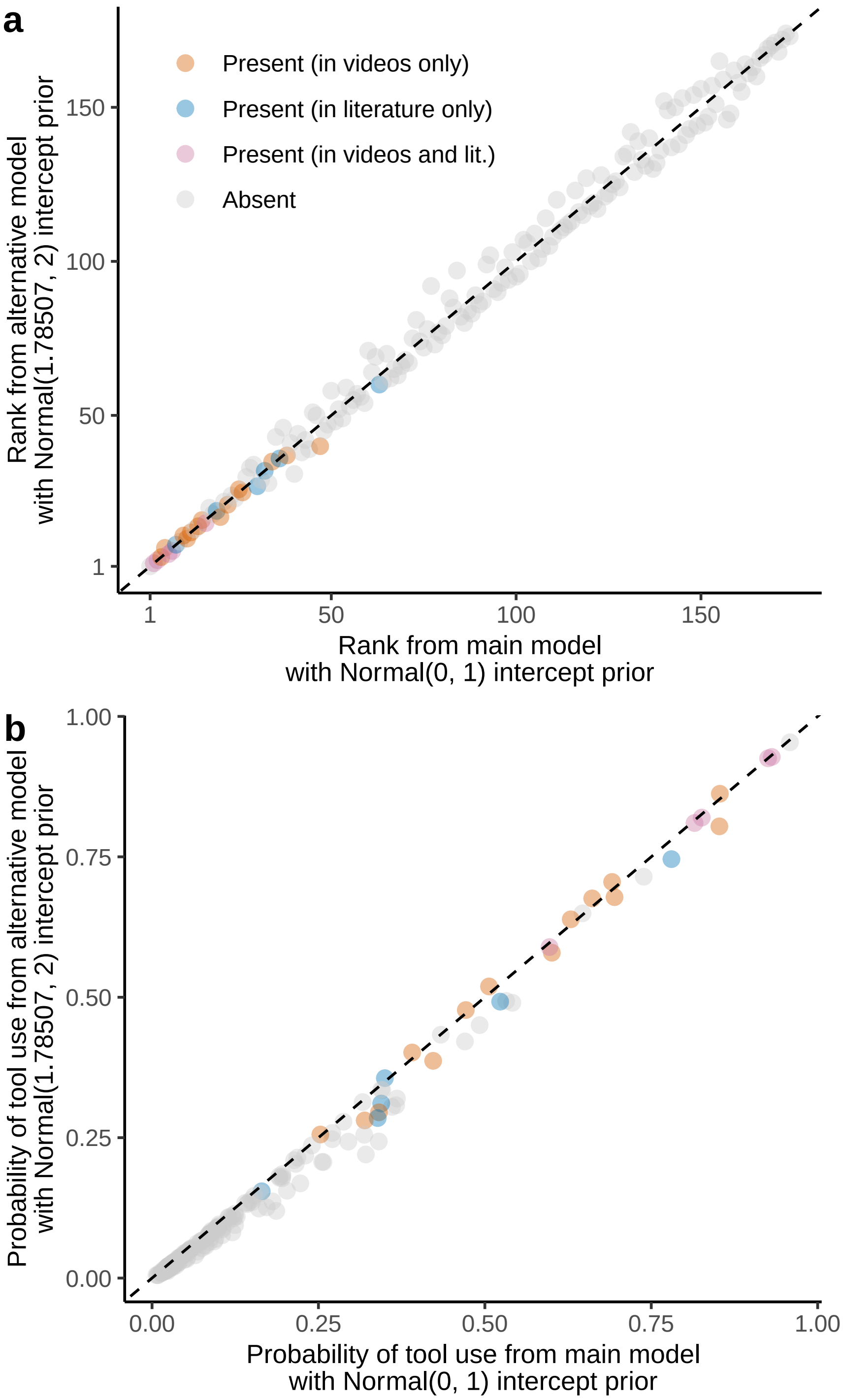


*Supplementary Figure 13*. *Results of sensitivity analysis.* The phylogenetic survival cure model was fitted with either a standard Normal(0, 1) prior on the intercept or an alternative Normal(1.78507, 2) prior on the intercept. This latter prior is wider on the logit scale and roughly converts to a 0.86 prior probability of non-tool-use (or a 0.14 prior probability of tool-use, which is the proportion of tool users in the dataset). The sensitivity analysis showed that changing this intercept prior did not have a marked impact on (a) the posterior rankings of parrot species from 1st to 174th or (b) the median posterior probabilities of tool use for parrot species.

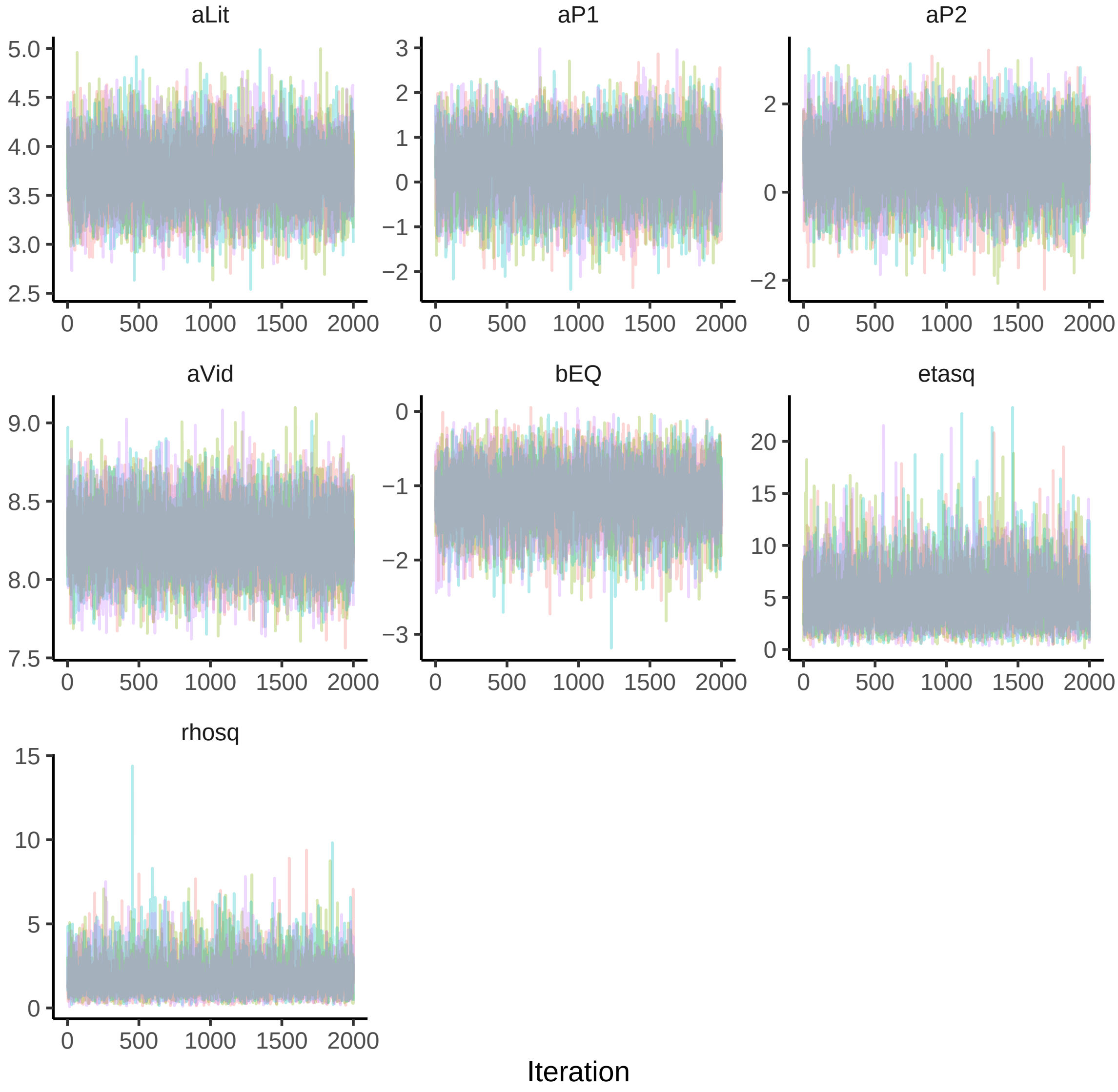


*Supplementary Figure 14. Trace plots for the Bayesian phylogenetic survival cure model.* Only four chains are shown for ease of presentation.

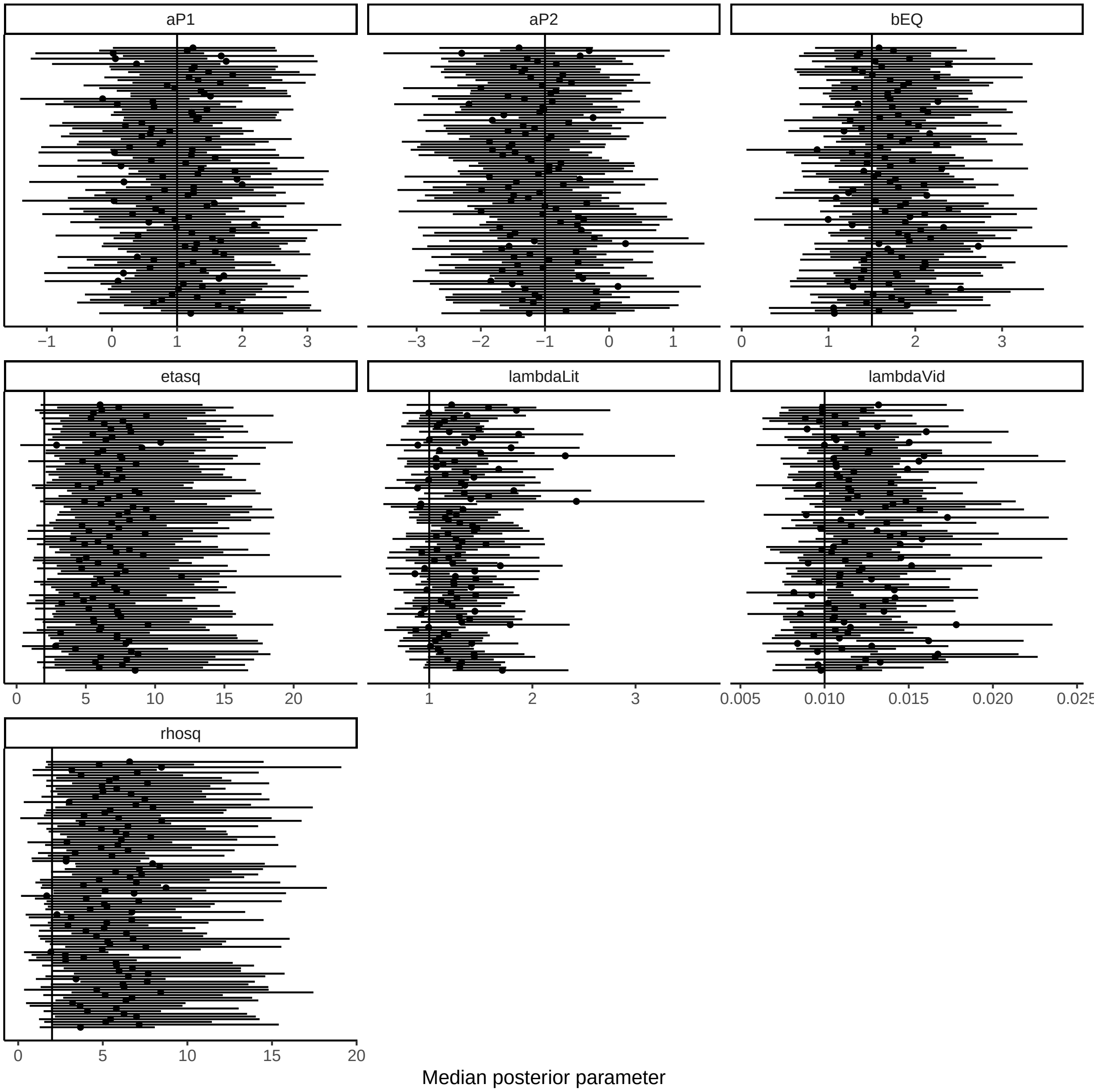


*Supplementary Figure 15. Posterior estimates from Bayesian survival cure models fitted to 100 datasets simulated with known parameters.* Each dataset consisted of 100 species. Known parameters are presented as solid vertical lines, whereas points and horizontal lines represent posterior medians and 95% credible intervals. The models were successfully able to recapture the parameters from the simulated datasets.
